# Syndecan is critical for *Drosophila* CNS and PNS glia function

**DOI:** 10.64898/2026.02.24.707778

**Authors:** Duo Cheng, Zhiheng Luo, Vanessa J Auld

## Abstract

Glia are irreplaceable components for the nervous system development and function. However, the cellular mechanisms each glial layer utilizes to communicate with each other and the extracellular environment is not well characterized. Here, we investigated the role of a heparan-sulfate proteoglycan, Syndecan (Sdc), in regulating glial cell function and development in the *Drosophila* nervous system. Sdc is expressed throughout multiple glial layers and loss of Sdc in all glia resulted in disruption of both central and peripheral glia. Within the CNS loss of Sdc in all glia lead to reduced brain lobes and disruption of neuroblast proliferation. In the PNS, loss of Sdc in different glial layers resulted in impaired ensheathment in wrapping glia and abnormal septate junction morphology in subperineurial glia. We focused on the outer layer of perineurial glia and found ensheathment defects and a reduction in glial numbers with Sdc loss. These phenotypes mirror those previously observed with the loss of integrins and a mutation in the integrin β-subunit enhanced the phenotypes observed with loss of Sdc within the perineurial. Thus, our results indicate Sdc has multiple roles in *Drosophila* nervous system development including as an integral component in regulating glial cell morphology, maintaining neuroblast populations within the optic lobe and in mediating glial-ECM interactions.

## Introduction

The ability of cells of all types to communicate among themselves and with the extracellular environment is a pivotal aspect of their overall functioning. As such, glia rely heavily on efficient adhesion and communication with their surroundings to ensheathe, protect and modulate the nervous system. However, the mechanisms by which glia communicate with each other and with the extracellular matrix is poorly understood.

The nervous system of Drosophila is encased in a robust extracellular matrix that covers both the central and peripheral nervous systems and underneath this matrix are a series of glial layers. Both the CNS and PNS are covered by a thin layer of surface glia that create the blood-brain and blood-nerve barrier and is created by the perineurial and subperineurial glia. The subperineurial are large polyploid cells, which establish a permeability barrier via the septate junctions, isolating the neurons from the circulating hemolymph filling the body cavity (Auld et al., 1995; Baumgartner et al., 1996; Stork et al., 2008). The perineurial glia lie between the ECM and the subperineurial glia. Perineurial glia are small diploid cells that are mobile and undergo mitosis throughout larval development, spreading across the surface of the entire nervous system by the late third instar stage (Awasaki et al., 2008; von Hilchen et al., 2013). To communicate with the overlying ECM, perineurial glia express transmembrane receptors and release metalloproteases, such as integrin and ADAMTS-A respectively, to direct CNS structural remodeling during development and structural integrity in the larval stages (Meyer et al., 2014; Skeath et al., 2017; Xie and Auld, 2011). Loss of components of the focal adhesion complex leads to incomplete ensheathment by the perineurial glia (Meyer et al., 2014; Xie and Auld, 2011). Though studies in both *Drosophila* and vertebrates have delved into the importance of integrin-mediated focal adhesion in glial-ECM adhesion, questions remain about the identities of other transmembrane receptor candidates besides integrin contribute to the cell-ECM adhesion in glia during development.

The syndecan family proteins constitute a group of transmembrane heparan sulfate proteoglycans (HSPGs) cell surface receptors that consist of a core protein with heparan sulfate glycosaminoglycan chains covalently attached to the ectodomain. There are four mammalian syndecans (syndecan-1, –2, –3, and –4) each having unique characteristic structural and expression profiles, though similarities do exist. Functional redundancy has led to challenges in studying the *in vivo* functions of mammalian Sdc (Couchman et al., 2015). In comparison, *Drosophila* contains a single *Sdc* gene. Drosophila Sdc is present in the embryonic nervous system, and in the both larval neuropile and neuromuscular junction in late larva stages (Spring et al., 1994; Stork et al., 2014; Nguyen et al., 2016). Yet, the presence and function of Sdc in glial populations remains poorly explored.

Mammalian syndecans are enriched in focal adhesion and growth-factor signaling complexes, thus placing syndecan in multiple signaling pathways and cellular functions. Syndecan acts in cooperation with integrin to direct focal adhesion and stress fiber formation in fibroblasts to mediate cell migration (Morgan et al., 2007). Additionally, syndecan acts to concentrate local ECM-ligand and growth factor levels, allowing these ligands to bind to their corresponding receptors with higher efficiency (Wu et al., 2003; Cheng et al., 2016). Sdc is important for a range of cell migration and adhesion pathways during *Drosophila* development, including axon migration and cardiogenesis (Johnson et al., 2004; Steigemann et al., 2004; Rawson et al., 2005; Chanana et al., 2009; Knox et al., 2011; Smart et al., 2011). Moreover, *Drosophila* Sdc plays a conserved role in trophic factor pathways such as FGF signaling, for example during tracheal cell migration (Lin et al., 1999; Schulz et al., 2011). Similarly, loss of HSPGs cause tracheal defects (Lin et al., 1999). In astrocyte-like glia, Sdc directs morphogenesis through FGF signaling and concentrates FGF ligands similar to vertebrate syndecan (Stork et al., 2014). However, the characterization of Sdc’s role in other Drosophila glia types has not been done.

To understand the role of Sdc in mediating glial development we analyzed the distribution and function of Sdc in larval glia. We observed Sdc is present in multiple glial layers in the third instar larvae, particularly prominent in the surface glial layers encasing the CNS, and all three glial layers in the PNS. When Sdc was knockdown in all glia, we observed significant changes in brain lobe size, elongation of the ventral nerve cord and disruption of animal locomotion and survival. When Sdc was reduced in specific glial layers several distinct phenotypes were observed within all three layers of the PNS. Loss of Sdc in wrapping glia disrupted axon ensheathment and loss of Sdc in the subperineurial glia led to localized swelling and disruption of septate junction protein distribution. Knockdown of Sdc in the perineurial glia resulted in incomplete ensheathment and migration defects along with reduced glial numbers. For these latter defects, we found that Sdc and integrin genetically interact to mediate perineurial glial ensheathment but not glial proliferation. Overall, we have identified Sdc as a key component in proper ensheathment and function of multiple glial cell types.

## Results

### Syndecan is expressed in a range of glia in the larval nervous system

To determine the expression pattern and protein localization of Sdc among the glial populations within *Drosophila* CNS and PNS, we used Sdc endogenously tagged with GFP (Sdc::GFP) (**Supplemental Fig. 1D**) (Buszczak et al., 2007). Animals homozygous for the *Sdc::GFP* allele displayed no gross nervous system defects and were viable. This suggests that the GFP insertion does not significantly impact Sdc function and localization and thereby is an accurate representation of normal Sdc expression. Sdc is expressed throughout the nervous system of 3rd instar larvae, including the brain lobes, the ventral nerve cord, and the peripheral nerves. To view the distribution of Sdc::GFP specifically in glia we used super-resolution imaging and marked all glial membranes using pan-glial driver *repo-GAL4* to express *mCD8::RFP* (**Fig. 1**). We found that Sdc is expressed in a punctate pattern that is often linear and outlines the edges of the glial membranes in the brain lobe (**Fig. 1A**), ventral nerve cord (**Fig. 1B**) and peripheral nerves (**Fig. 1C**). Additionally, we observed robust Sdc::GFP outlining cell boundaries within the brain lobe and ventral nerve cord that did not colocalize with any glial cells (**Fig. 1A,1B**, yellow arrow) likely due to expression in neurons, consistent with previous studies (Johnson et al., 2004; Steigemann et al., 2004). Moreover, Sdc puncta localized to the superficial glial layers, suggesting that Sdc is expressed in the perineurial glia population (**Fig. 1A-C**, white arrow). The localization of Sdc was further confirmed using a second independent GFP insertion in the *Sdc* gene (Sdc::GFP.M) (Nagarkar-Jaiswal et al., 2015) (**Fig. 1D**).

**Figure 1.**
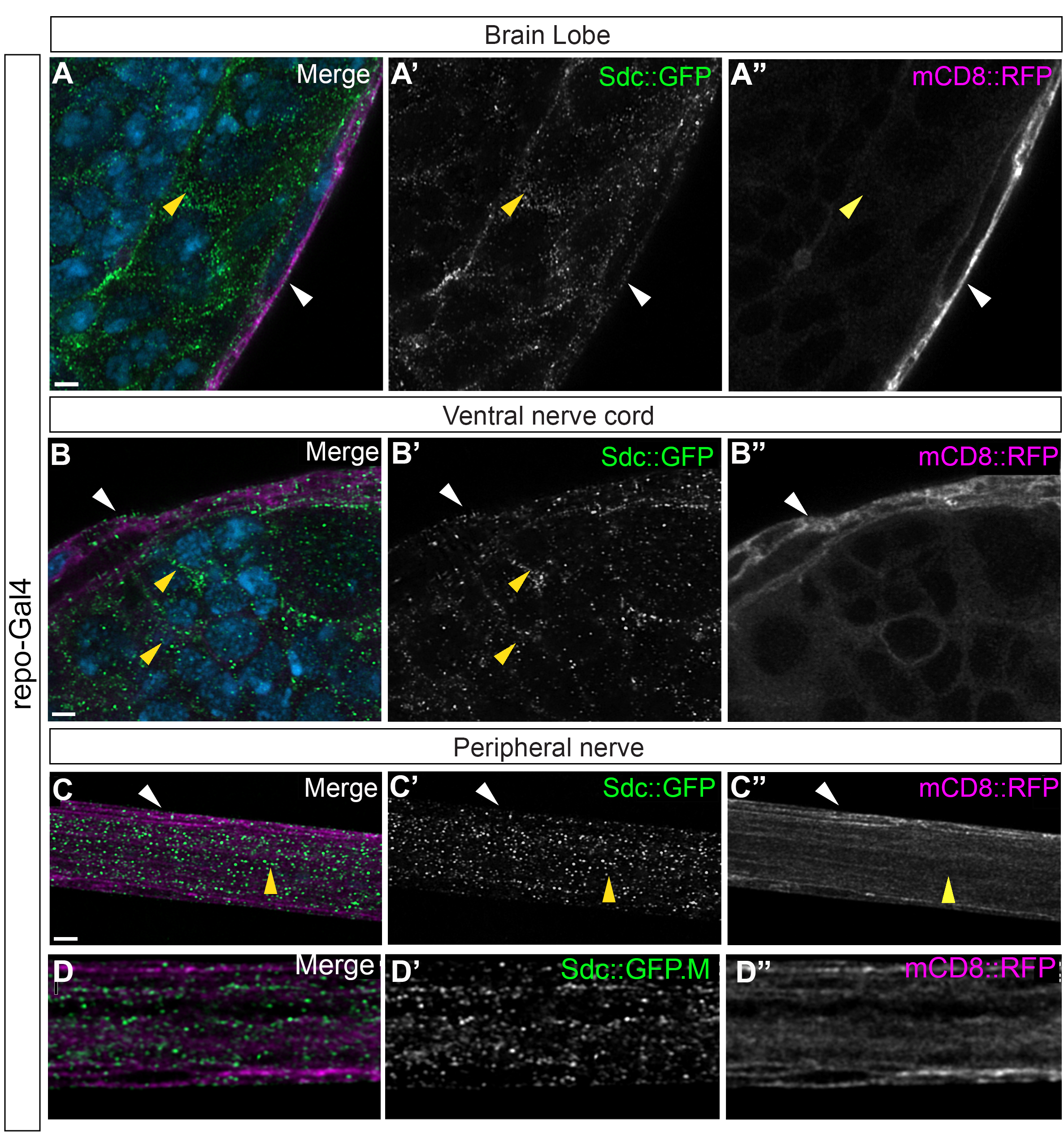
Sdc is expressed by glia in the central and peripheral nervous systems. Super resolution images of the brain lobe (**A**), ventral nerve cord (**B**) and peripheral nerves (**C-D**) of third instar *Drosophila* larvae expressing endogenously tagged Sdc::GFP (green). Glial cell membranes are labelled using mCD8::RFP (magneta) driven by the pan-glia driver repo-Gal4. Cell nuclei are labelled with DAPI (blue). Scale bar, 2 µm (A) Sdc::GFP is found throughout the brain lobe including deep (yellow arrowheads) and within the superficial perineurial and subperineurial glial layers (white arrowhead). (B) Sdc::GFP is found throughout the ventral nerve cord (yellow arrowheads) including within the superficial perineurial and subperineurial glial layers (white arrowhead). (C) Sdc::GFP is found through the peripheral nerve in all three glial layers including the superficial perineurial glia (white arrowhead) and deep within the nerve (yellow arrowhead) either in the wrapping glia or peripheral axons. (D) Sdc:: GFP.M (green) visualized using an anti-GFP antibody had a similar distribution to Sdc::GFP with puncta throughout the glial membranes.

To better characterize Sdc expression within specific glial layers, we assessed Sdc::GFP localization specifically within individual glial layers of the CNS and PNS. For our analysis in the CNS, we used a membrane-bound RFP (mCD8::RFP) paired with Gal4 drivers to the most superficial glial layers that surround the CNS and PNS, specifically the perineurial glia using *46F-Gal4* (Xie and Auld, 2011) and the subperineurial glia using the *SPG-Gal4* driver (Stork et al., 2008). Within the CNS (**Fig. 2A-B**), Sdc::GFP puncta are closely associated with the perineurial glial membrane at the glial-ECM interface (**Fig. 2A**). There were also instances where Sdc puncta localized to the perineurial-subperineurial cell-cell interface (**Fig. 2B**, yellow arrow) and patches of Sdc puncta localized to the subperineurial glial membrane (**Fig. 2B**). Within the peripheral nerve Sdc was observed within all three glial layers (**Fig. 2C-E; yellow arrows**), with puncta found within the perineurial glia (**Fig. 2C**), the subperineurial glia (**Fig. 2D**) and the wrapping glia (**Fig. 2E**). However, it is likely that Sdc is also expressed in the peripheral axons (Smart et al., 2011).

**Figure 2.**
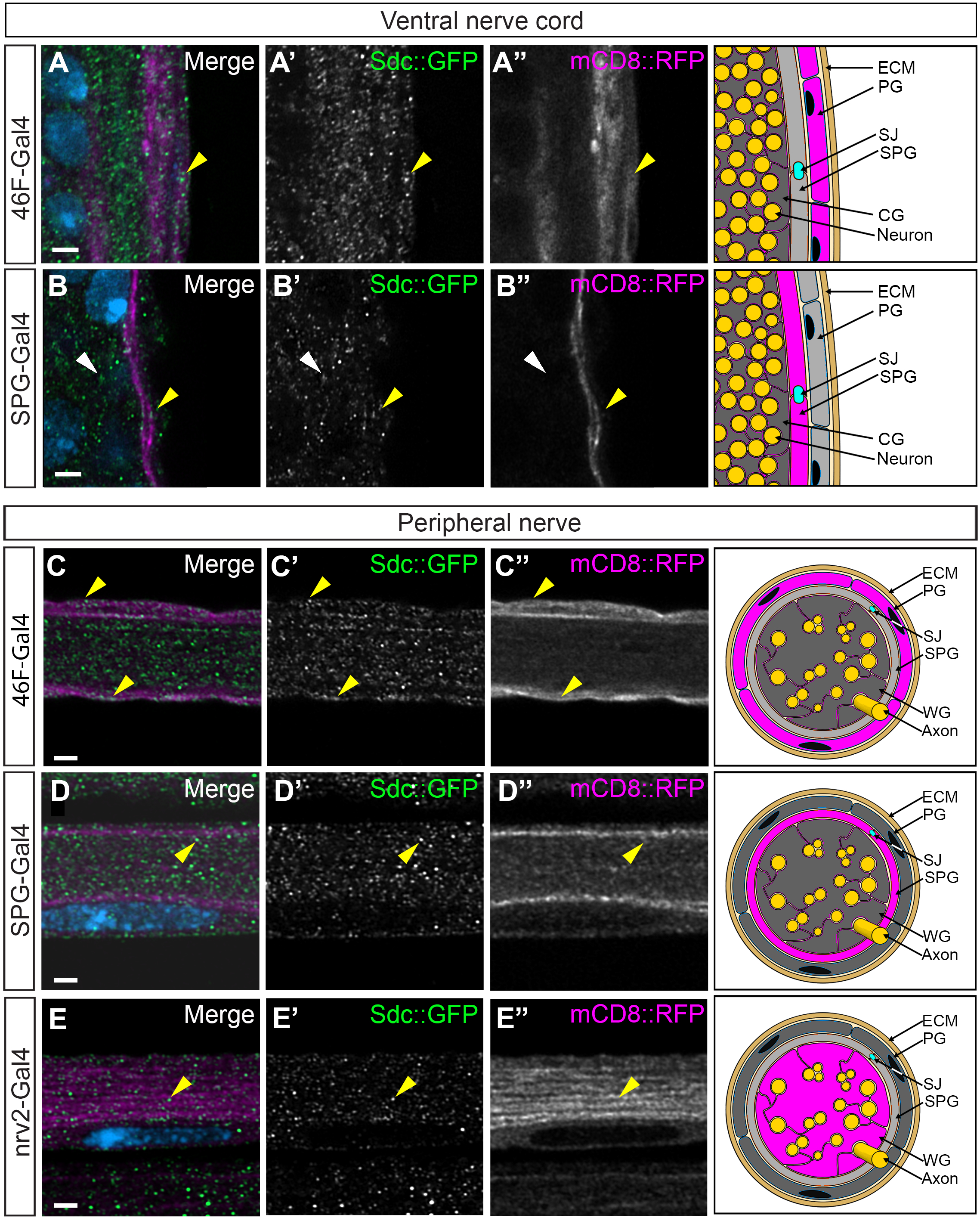
Sdc is expressed in multiple glial layers. Third instar *Drosophila* larvae expressing endogenously tagged Sdc::GFP (green) with distinct glial layer membranes labelled using specific Gal4 lines driving mCD8::RFP (magneta). Cell nuclei are labelled with DAPI (blue). Scale bar, 2 µm. Cartoons to the right depict the labeled glia (magenta) – ECM: extracellular matrix. PG: Perineurial glia. SJ: septate junction. SPG: subperineurial glia. CG: cortex glia. WG: wrapping glia. (**A-B**) Sdc::GFP is found in glial layers surrounding the CNS. (**A**) Perineurial glia (46F-Gal4) on the outer layer of the VNC and (**B**) subperineurial glia (SPG-Gal4) flanking the perineurial glia express Sdc (yellow arrows). Sdc is also expressed in deeper glial layers including the cortex glia (CG, white arrowheads) which surround the neuronal cell bodies in this region. (**C-E**) Sdc::GFP is found in all three glial layers (yellow arrows). (**C**) Perineurial glia (46F-Gal4) surround each peripheral nerve, (**D**) subperineurial glia (SPG-Gal4) flank the perineurial glia and (**E**) wrapping glia (Nrv2-Gal4) wrap peripheral axons.

### Glial Syndecan is required for survival

Given the strong expression of Sdc within both CNS and PNS glia, our goal was to establish the overall requirement of Sdc. To do so, we reduced Sdc expression using verified RNAi lines Sdc RNAi-1 (GD4545) and Sdc RNAi-2 (KK108799) (Knox et al., 2011; Nguyen et al., 2016; Eldridge-Thomas et al., 2025) that target different regions of the *Sdc* transcript (**Supplemental Fig. 1D**). We used the pan glial driver, *repo-GAL4,* and *UAS-Dicer2 (Dcr2)* to enhance RNAi efficiency along with the membrane marker, mCD8::RFP. Expression of the two independent RNAi lines lead to the loss of Sdc::GFP within the glial layers (**Supplemental Fig. 1 A-C”**).

We tested adult viability after Sdc knockdown in all glia and found strong effects on adult viability where on average 0% of Sdc RNAi-1 (*repo>Dcr2,Sdc RNAi-1*) and 21% of Sdc RNAi-2 (*repo>Dcr2,Sdc RNAi-2*) pupae eclosed compared to 100% for controls (*repo>Dcr2)* (**Supplemental Fig. 1E**). To test the specificity of the glial knockdown, we also tested the viability of *Sdc* mutants using a heteroallelic combination of *Sdc* loss of function mutants: a deficiency that removed the *Sdc* gene (*Df(2R)48, ubi-Sara)* (Johnson et al., 2004) and the *Sdc^97^* allele (a deletion that removes the transcription start site, the first exon and part of the first intron (Steigeman et al., 2004)) (**Supplemental Fig. 1D**). Both *w^1118^* and the heterozygous deficiency (*Df(2R)48, ubi-Sara*/+) had on average 100% survival, while the allelic combination (*Df(2R)48, ubi-Sara/Sdc^97^)* had, on average, 43% survival (**Supplemental Fig. 1E**). It should be noted that the knockdown of Sdc using RNAi in any of the specific glial layers (perineurial, subperineurial and wrapping glia) did not impact survivability as all progeny (100%) survived to adult stages. Overall, our results suggest that glial Sdc is required for survival and that Sdc-RNAi-1 generates the most effective knockdown (similar to prior observations (Eldridge-Thomas et al., 2025)).

### Glial Syndecan is important for animal locomotion

Given the strong expression in both CNS and peripheral glia, we next tested the effect of Sdc knockdown on locomotion in all glial or specific glial layers including the perineurial, subperineurial and wrapping glia. The knockdown of Sdc in all glia (*repo>Dcr2, Sdc RNAi-1*; *repo>Dcr2, Sdc RNAi-2*) lead to a significant reduction in overall movement compared to control (*repo>Dcr2)* (**Fig. 3A-C**). Further quantification of the average speed (**Fig. 3D**), and total distance travelled (**Fig. 3E**) revealed the impaired locomotion associated with *RNAi-1* was the most severe, showing a more than 50% reduction in average speed and distance traveled compared to control. A significant, but less severe decrease in movement was observed with Sdc RNAi-2. We next tested the loss of Sdc in the subperineurial glia that create the blood-brain barrier and blood-nerve barriers in the CNS and PNS respectively. Using a SPG specific driver, we saw a reduction in animal movement across both RNAi groups (*SPG>Dcr2, Sdc RNAi-1*; *SPG>Dcr2, Sdc RNAi-2*) compared to control (*SPG>Dcr2*) (**Fig. 3F,G**), though the effect was not a strong as with the pan-glial driver. Conversely, when knock down was assessed in the wrapping glia, only the stronger RNAi-1 (*Nrv2>Dcr2, Sdc RNAi-1*) significantly affected larval locomotion in comparison to RNAi-2 (*Nrv2>Dcr2, Sdc RNAi-2)* (**Fig. 3H,I**). Surprisingly, the loss of Sdc in the perineurial glia with the stronger *RNAi-1* (*46F>Dcr2, Sdc RNAi-1*) did not disrupt animal locomotion compared to control (*46F>Dcr2*) (**Fig. 3J)** but there was a significant increase in both distance travel and speed with *RNAi-2* (*46F>Dcr2, Sdc RNAi-2)* (**Fig. 3K**) after perineurial glial knockdown. Overall, the loss of Sdc in all glia or a subset of glia had a strong effect on larval locomotion.

**Figure 3.**
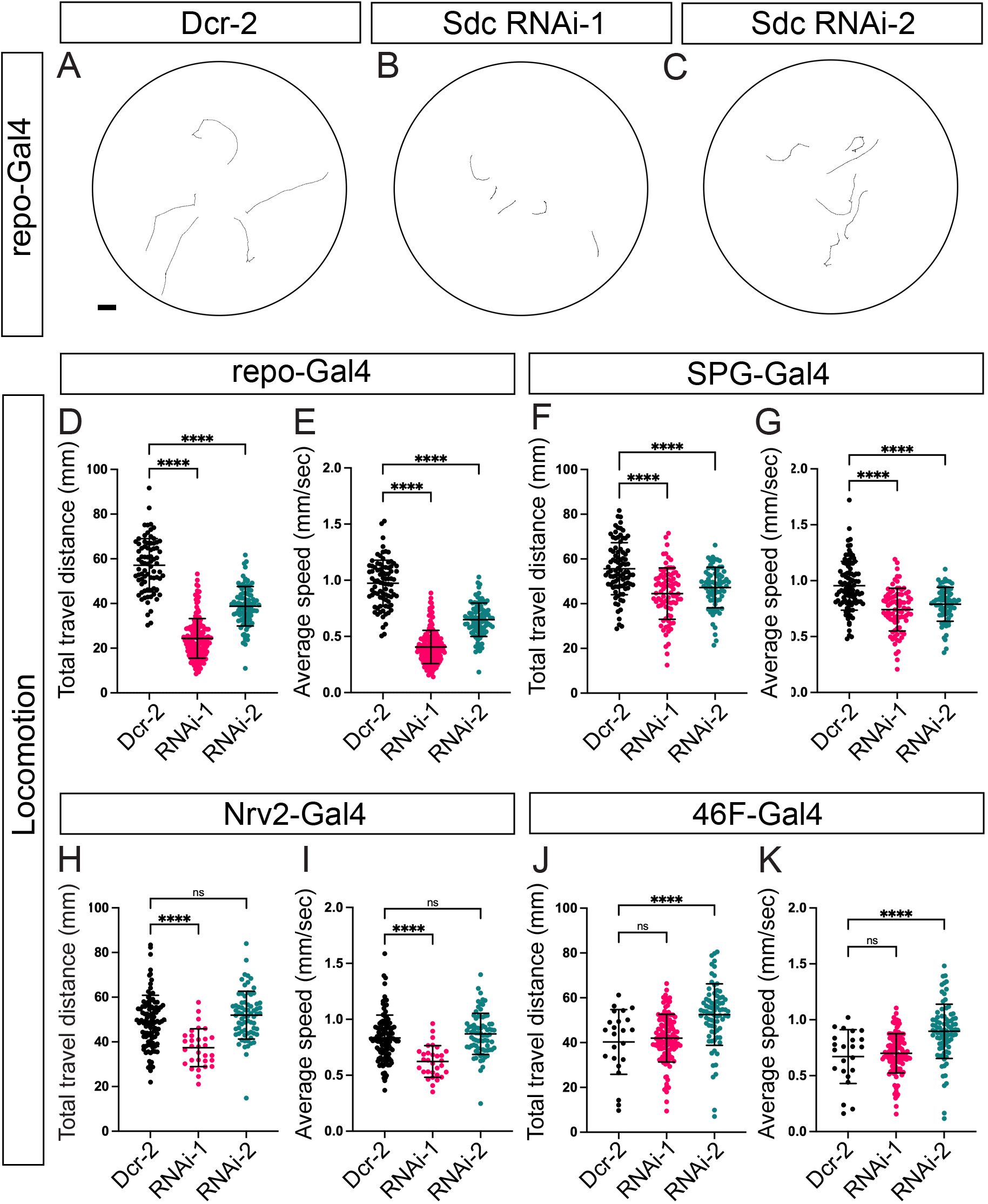
Sdc is required for proper locomotion. **(A-C)** Representative movement trajectory of control (Dcr-2) (**A**), Sdc RNAi-1+Dcr-2 (**B**), Sdc RNAi-2+Dcr-2 (**C**), expressed using the pan-glial driver, repo-GAL4. Each line is indicative of one larva’s movement in 60 seconds with the starting point near the center. Scale bar, 10mm. (**D-K**) Knockdown of Sdc using two independent RNAi lines compared to Dicer-2 (Dcr2) alone driven with specific glial drivers. All RNAi lines were co-expressed with Dicer-2. Each analysis was a One-way ANOVA, Dunnet’s post-hoc multiple comparison. **** p<0.0001; *** p<0.001; ns – not significant. **(D-E)** All glia (repo-Gal4): Knockdown in all glia significantly reduced total travel distance (**E**) and average larval speed (**F**) for both Sdc RNAi-1 and Sdc RNAi-2 compared with control (Dcr2). (**F-G**) Subperineurial glia (SPG-Gal4): Knockdown in all glia significantly reduced total travel distance (**F**) and average larval speed (**G**) for both Sdc RNAi-1 and Sdc RNAi-2 compared with control (Dcr-2). **(H-I)** Wrapping glia (Nrv2-Gal4): Knockdown of Sdc in wrapping glia significantly decreased total travel distance (H) and average larval speed (I) for Sdc RNAi-1 when compared to control. No significant reduction was observed with Sdc RNAi-2. **(J-K)** Perineurial glia (46F-GAL4): Knockdown of Sdc in perineurial glia significantly increased total travel distance (J) and average larval speed (K) with Sdc RNA-2 compared to control. Sdc RNAi-1 had no significant differences to Dcr-2 control.

### Glial Syndecan is necessary for CNS morphology

Given the strong effects of pan-glial knockdown on locomotion, our next assessment was to visualize any changes to CNS and glia morphology using a membrane marker (mCD8::GFP) in conjunction with Sdc RNAi and the pan glial driver, *repo-Gal4*. In comparison to control animals (*repo>Dcr-2, mCD8::GFP*) (**Fig. 4A**), we found various morphological defects spanning the CNS upon Sdc knockdown. The VNC appeared elongated (**Fig. 4B,C**, white arrows), and the brain lobes were noticeably smaller (**Fig. 4B,C**, yellow arrows). We quantified these effects and found a statistically significant increase in the ratio of VNC to body length with both RNAi lines (*repo>Dcr-2, mCD8::GFP, Sdc RNAi-1*; *repo>Dcr-2, mCD8::GFP, Sdc RNAi-2)* (**Fig. 4D**). Compared to control (*repo>Dcr-2, mCD8::GFP)*, where the VNC is on average 9.8% of the body length, we saw an increase where the VNC extends to an average of 11% and 10.8% of the body length in *RNAi-1* and *RNAi-2*, respectively. We also quantified the severity of the brain lobe size reduction (**Fig. 4E**). By measuring the brain lobe at the widest point (from outer edge to center), we saw a 27.8% decrease in brain lobe width with *RNAi-1* and a 13.4% reduction in *RNAi-2* compared to control.

**Figure 4.**
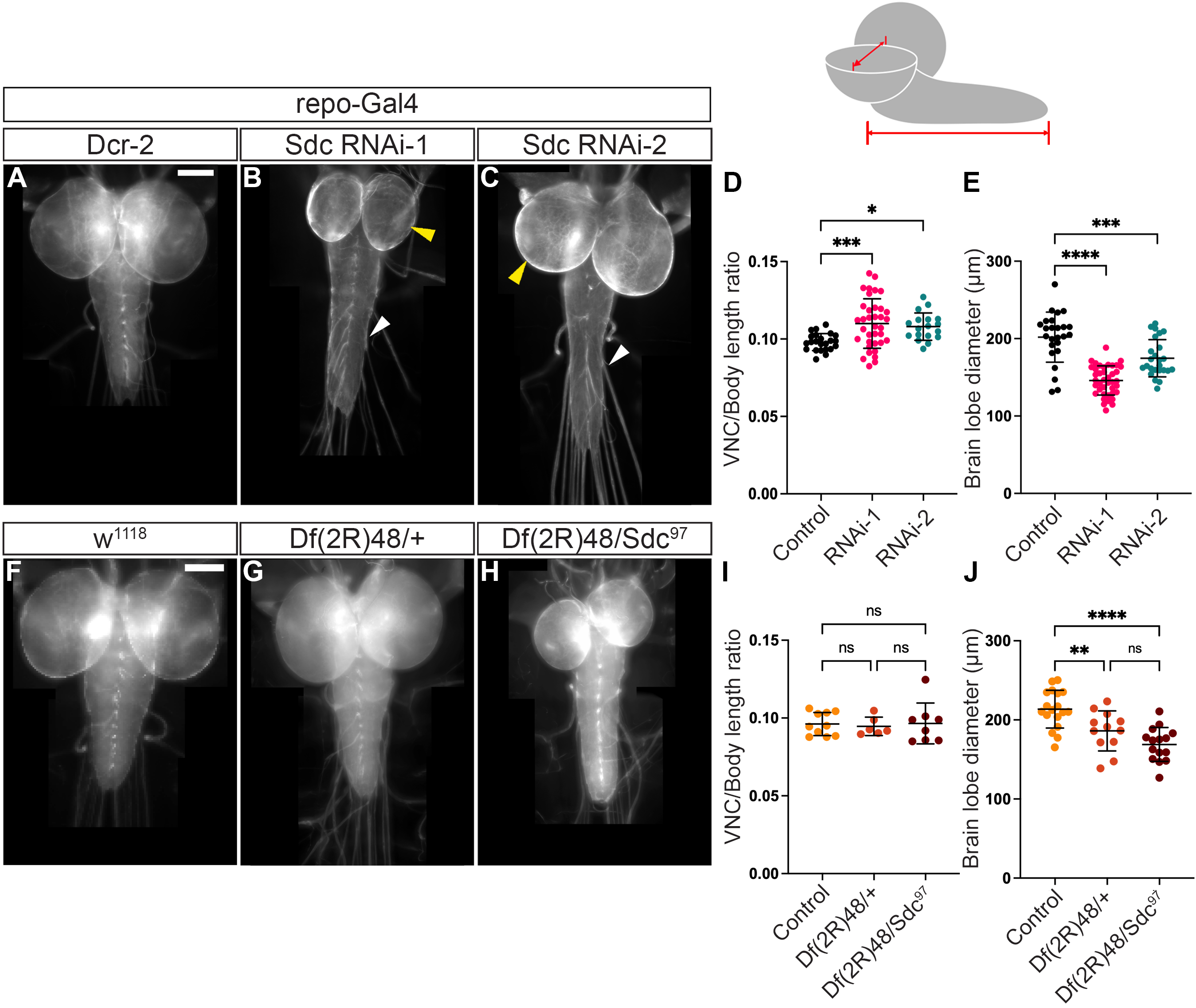
Loss of Sdc in all glia leads to CNS defects. Comparison of loss of Sdc in all glia using the pan-glial driver (repo-Gal4) with Sdc-RNAi (**A-E**) or Sdc loss of function mutants (**F-J**). Glial membranes were labeled with repo-GAL4 driving the expression of mCD8::GFP (gray). The cartoon depicts the larval CNS with the dimensions measured (red lines) for the brain lobe diameter (**E,J**) and the VNC length (**D,I**). All RNAi lines were co-expressed with Dicer-2. All graphs represent a One-way ANOVA, Dunnett multiple comparison test: **** p<0.0001; *** p<0.001; **p<0.01; *p<0.05; ns – not significant. Scale bar, 100 µm. **(A**-**C)** Representative images of the brain lobes and VNCs of third instar *Drosophila* larvae of control Dcr-2 alone (**A**), Sdc RNAi-1 (**B**), and Sdc RNAi-2 (**C**). Knockdown of Sdc results in length and morphological defects in the ventral nerve cord (white arrows, **B**-**C**) and reduced brain lobes (yellow arrows, **B-C**) compared to control (**A**). **(D)** Pan-glial Sdc RNAi knockdown resulted in elongation of the VNC compared to control (Dcr-2) when measuring VNC length as a ratio of the larval body length. **(E)** Pan-glial Sdc RNAi knockdown resulted in smaller brain lobes compared to control (Dcr-2). (**F-H)**, Representative images of the third instar larvae CNS of control (*w^1118^*) (**F**), Sdc heterozygous mutant (*Df(2R)48, ubi-Sara/+)* (**G**), Sdc homozygous mutant *Df(2R)48, ubi-Sara/Sdc^97^* (**H**). **(I)** *Sdc* homozygous mutants (*Df(2R)48, ubi-Sara/Sdc^97^*) and *Sdc* heterozygous mutant (*Df(2R)48, ubi-Sara/+*) did not have a significant elongation of the VNC as a ratio of larval body length when compared to control (*w^1118^*). **(J)** *Sdc* homozygous (*Df(2R)48, ubi-Sara/Sdc^97^*) and heterozygous mutants (*Df(2R)48, ubi-Sara/+*) had significantly smaller brain lobes when compared to control (*w^1118^*).

We next confirmed these phenotypes using the heteroallelic combination of *Sdc* loss of function mutants using *repo-Gal4* with mCD8::GFP to label the glial membranes. We observed no significant differences in the VNC to body length ratio with the heterozygous *Df(2R)48, ubi-Sara/+* or the heteroallelic combination of *Df(2R)48, ubi-Sara/Sdc^97^* compared to control (*w^1118^*) (**Fig. 4F-I**). However, the brain lobes were significantly reduced in both the heterozygous *Df(2R)48, ubi-Sara/+* and the heteroallelic combination of *Df(2R)48, ubi-Sara/Sdc^97^* compared to control (*w^1118^*) (**Fig. 4H**, yellow arrows; **Fig. 4J**). Overall, our results suggest that glial Sdc is required for CNS morphology and in particular regulates the size of the brain lobes.

### Glial Syndecan is needed for neuroblast proliferation in the CNS

Neurons account for the majority of the tissue volume in 3^rd^ instar larval brain lobes, with discrete regions of neuroblast proliferation that undergo cell division to generate new neurons (Egger et al., 2008). The brain lobe shrinkage we identified with pan-glial expression of the Sdc-RNAi lines could be associated with decreased proliferation or an increase in programmed cell death. Thus, we tested the effects that pan-glial loss of Sdc has on neuroblasts in the larval brain lobe (**Fig. 5**). We used phosphohistone-3 (pHis-3) as a marker for mitotic events in the brain lobe. In controls (*repo>Dcr2, mCD8::GFP)*, we observed a zone of active proliferation, as marked by the robust signal of pHis-3 surrounding by dense-area of nuclei marked by DAPI (**Fig. 5A-A’’’**). This region corresponds to the optic lobe outer proliferation zone (Egger et al., 2007) and is robustly labeled using the neuroblast marker Deadpan (Dpn) (San-Juán and Baonza, 2011) (**Fig. 5D-D’’’**). Knockdown of Sdc in all glia lead to a reduced size of the brain lobe and the outer proliferation zone (**Fig. 5B-C’’’; E-F’’’**). *RNAi-1* demonstrated the most severe phenotype, where a large portion of the outer proliferation zone (white dash lines) was greatly reduced. With RNAi-2, we observed a decrease in the neuroblast region, though it was not as dramatic as RNAi-1. When quantified, we observed a significant reduction of active cell divisions (pHis-3) within the neuroblasts with RNAi-1 (*repo>Dcr2, Sdc RNAi-1*) to 70% of control but not RNAi-2 (*repo>Dcr2, Sdc RNAi-2)* at 94% compared to control (*repo >Dcr2*) (**Fig. 5G**). Similarly, when we quantified the volume of the neuroepithelia region within the optic lobe there was a significant loss in size with the RNAi-1 knockdown (76%) compared to control (**Fig. 5H**). While RNAi-2 lead to smaller volume on average compared to control (88%), this was not a statistically significant difference.

**Figure 5.**
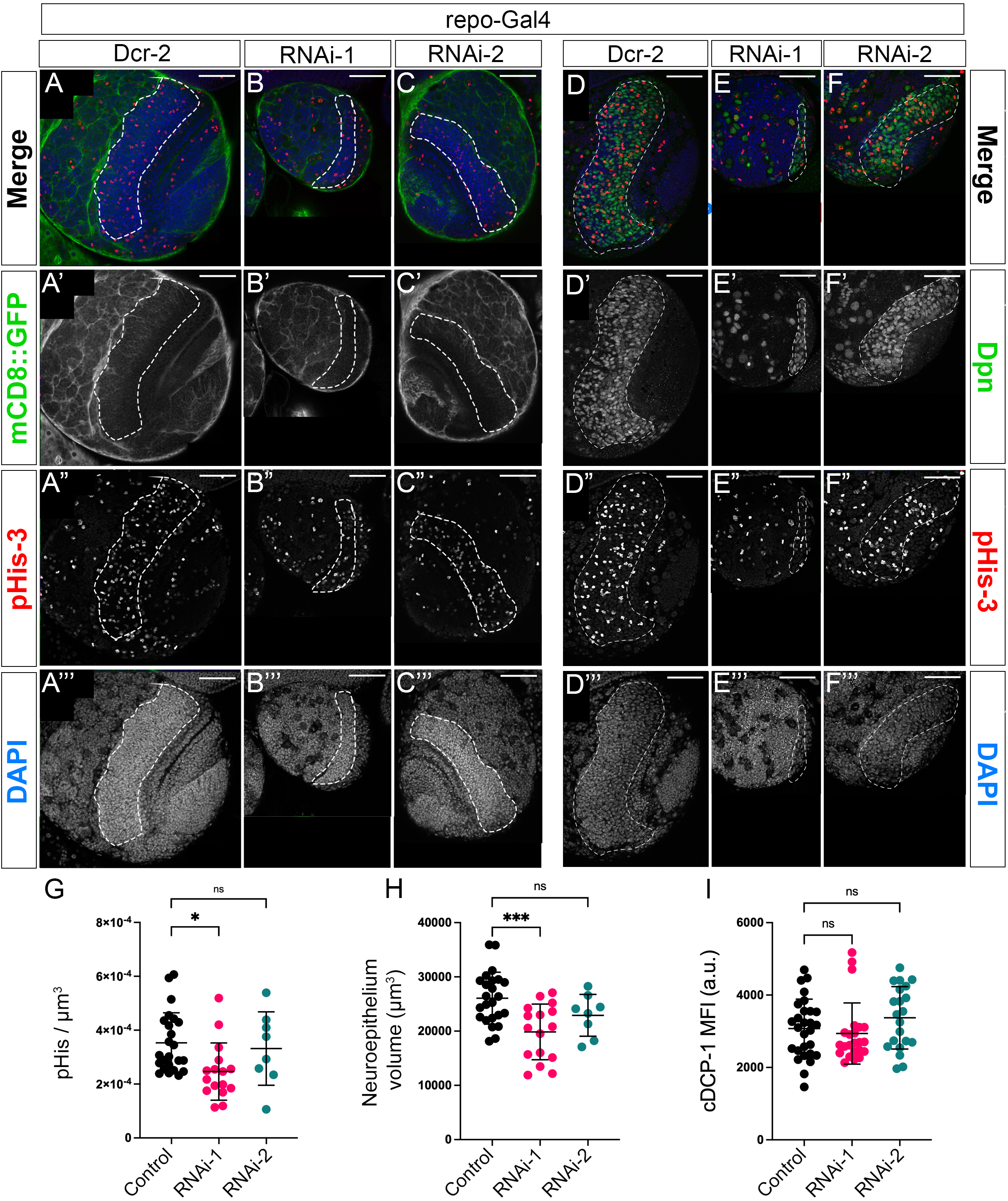
Neuroblast numbers and proliferation requires glial Syndecan. Representative images of the cross-sections of 3rd instar larval brain lobes with glial membranes labeled (**A-C**) or neuroblasts labelled (**D-F**). Mitotic cells were immunolabeled with anti-phosphohistone-3 (pHis-3, red). DAPI (blue) was used to mark nuclei and the cell-dense outer proliferation belt within the optic lobe (highlighted with white dashed line). All RNAi lines were co-expressed with Dicer-2 and driven with repo-Gal4. All graphs represent a One-way ANOVA, Dunnett multiple comparison test: *** p<0.001; *p<0.05; ns – not significant. Scale bars, 50 µm (**A-C**) All glial membranes were labelled with membrane-bound GFP (mCD8::GFP, green) expressed by *repo-GAL4*. Notice the changes in brain lobe size, proliferation zone (white dashed line) and reduced pHis-3 staining within Sdc-RNAi-1(**B**) and Sdc-RNAi-2 (**C**) compared to control (Dcr2) (**A**). (**D-F**) Representative cross-section images showing the proliferation zone (white dashed line) with the neuroblast population identified using an anti-deadpan (Dpn, green) antibody, mitotic cells with pHis-3 (red) and all nuclei with DAPI (blue). Note the less abundant neuroblast and miotic events with SdcRNAi-1 (**E**) and Sdc RNAi-2 (**F**) compared to control (Dcr2) (**D**). (G) Quantification of the mitotic cells within the proliferation zone, with significantly less pHis3 positive cells with Sdc RNAi-1 but not Sdc RNAi-2 compared to control (Dcr2) (H) Quantification of the volume of the proliferation zone, with significantly less neuroepithelial volume with Sdc RNAi-1 and Sdc RNAi-2 compared to control (Dcr2). (I) An anti-cleaved *Drosophila* caspase-3 (cDCP-1) antibody was used to detect cell apoptosis. There was no observed increase in cDCP-1 within Sdc RNAi-1 and Sdc RNAi-2 compared to control (Dcr2).

To test if the reduction in the proliferation zone is due to increased apoptosis, we examined the activity of cleaved Death caspase-1 (cDCP-1), an effector protein in the apoptosis pathway (Song et al., 1997). In both RNAi groups, we did not observe a noticeable increase in the amount of cDCP-1 activity comparing to the control (**Fig. 5I**). This result suggests that the decrease in brain lobe size is due to a lack of proliferation and not an increase in cell death. These results indicated that glial Sdc is necessary for neuroblast proliferation and thus regulates brain lobe size.

### Sdc in the peripheral nerve is necessary for all three PNS glial layers

We next tested the effects that loss of Sdc had on peripheral nerves. The larval peripheral axons are encapsulated by three glial layers including the outermost perineurial glia, the blood-nerve barrier forming subperineurial glia and the inner most wrapping glia that ensheathe both motor and sensory axons. After pan-glial expression of Sdc RNAi (**Fig. 6B-C**”), the peripheral nerves frequently displayed swelling and glial membrane fragmentation with both RNAis (yellow arrows) compared to the compact and smooth layers in control (*repo >Dcr2, mCD8::GFP)* (**Fig. 6A-A”**). Quantification revealed that (**Fig. 6D**) both RNAis lines had significantly greater numbers of affected nerves compared to control where RNAi-1 (*repo>Dcr2, mCD8::GFP, RNAi-1*) had 100% of nerves affected and RNAi-2 (*repo>Dcr2, mCD8::GFP, RNAi-2)* had 14% of nerves.

**Figure 6:**
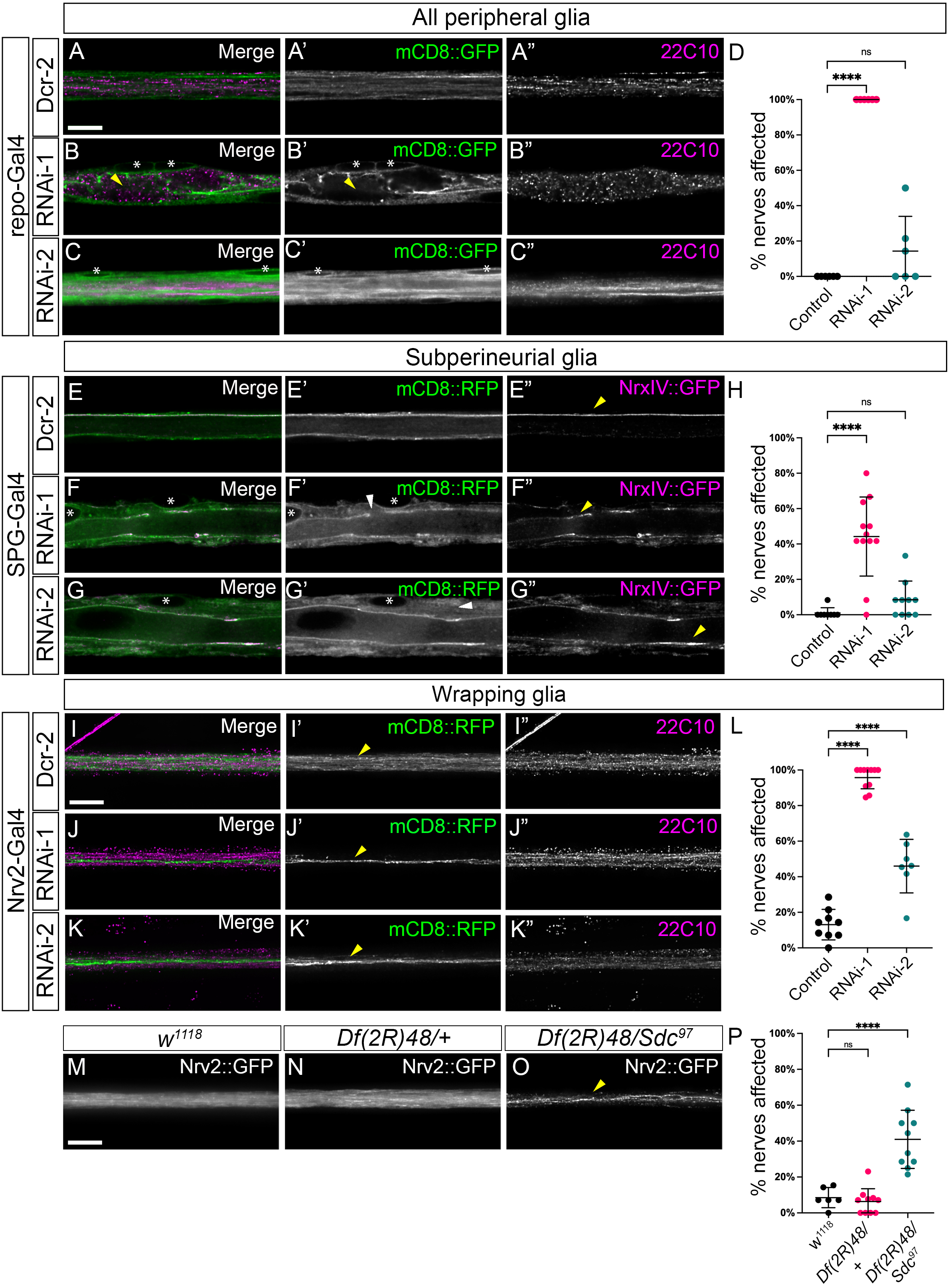
Knockdown of Syndecan disrupts peripheral glia morphology. Loss of Syndecan in all peripheral glia (**A-D**), the subperineurial glia (**E-H**) or the wrapping glia (**I-P**). All RNAi lines were co-expressed with Dicer-2. All graphs represent a One-way ANOVA, Dunnett multiple comparison test: **** p<0.0001; ns – not significant. The results are presented as means ± SD. ****p < 0.0001, ***p <0.001, *p < 0.05, n.s., not significant. Scale bar, 10 µm (**A-D**) Loss of Sdc in all glia. (**A-C**) Representative longitudinal-section images of all glial layers in third instar larvae peripheral nerve of control, Sdc RNAi-1, and Sdc RNAi-2. The pan-glial driver, repo-GAL4 was used to drive the expression of mCD8::GFP (green) to label all glial membranes, and axons immunolabeled for futsch (22C10, magenta). Sdc knockdown from RNAi results in the disruption of peripheral nerve integrity (**B**) when compared to control (**A**). Notice the swelling and absent axonal labelling within the peripheral nerve in Sdc RNAi-1(**B,** yellow arrow). Asterisks denote the position of nuclei. (**D)** The percentage of nerves showing morphological defects in the peripheral nerves is significant with Sdc RNAi-1 but not in Sdc RNAi-2 when compared to control (Dcr2). (**E-H**) Loss of Sdc in the subperineurial glia. (**E-G**) Representative longitudinal-section images of subperineurial glia in the third instar peripheral nerve of control (**E**), Sdc RNAi-1 (**F**), and Sdc RNAi-2 (**G**) expressing endogenously tagged NrxIV::GFP (magenta), a core septate junction protein. Subperineurial glia membranes were labelled using SPG-Gal4 driving the membrane marker mCD8::RFP (green). Septate junctions form a tight and continuous strand along the length of the nerve (**E”,** yellow arrowhead). In regions of glial swelling, Nrx-IV was mislocalized (**F”, G”,** yellow arrowheads) along with disorganized subperineurial membranes (**F’, G’**, white arrowheads). This is observed in all RNAi groups. Asterisks denote the position of nuclei. (**H**) The percentage of nerves showing subperineurial glia defects was significantly increased with Sdc RNAi-1 but not Sdc RNAi-2 when compared to control (Dcr2). (**I-P)** Loss of Sdc in the wrapping glia. (**I-K**) Representative longitudinal-section images of wrapping glia in third instar peripheral nerve of control (**I**), Sdc RNAi-1 (**J**) and Sdc RNAi-2 (**K**). Wrapping glia membrane is labelled using Nrv2-GAL4 driving mCD8::RFP (green) and axons immunolabelled with anti-futsch (22C10, magenta). In Sdc knockdowns, wrapping glia show a lack of radial ensheathment (**J’, K’,** yellow arrowheads) in contrast to control (**I’**). Axonal morphology appears unaffected by Sdc knockdown. (**L**) The percentage of nerves showing wrapping glia radial ensheathment defect in the peripheral nerve was significant for both Sdc RNAi (Sdc RNAi-1; Sdc RNAi-2) when compared to control (Dcr2). **(M-O**) Representative longitudinal-section images of wrapping glia in third instar larvae peripheral nerve of control (*w^1118^*) (**M**), Sdc heterozygous mutants (*Df(2R)48, ubi-Sara/+)* (**N**), Sdc homozygous mutants *Df(2R)48, ubi-Sara/Sdc^97^* (**O**). Wrapping glia membrane were labelled with Nrv2::GFP. In the *Sdc* null background, wrapping glia have decreased ensheathment (**O**, yellow arrowhead). (**P**) *Sdc* homozygous mutant (*Df(2R)48, ubi-Sara/Sdc^97^*) had significantly more ensheathment defects when compared to *Sdc* heterozygous mutant (*Df(2R)48, ubi-Sara/+*) and control (*w^1118^*).

As Sdc::GFP was localized to the subperineurial glial membrane, we assessed whether the morphology of subperineurial glia was affected with Sdc knockdown. We visualized glial morphology using mCD8::RFP under the control of *SPG-GAL4* and the septate junctions using Neurexin-IV, a core septate junction component endogenously tagged with GFP (*NrxIV::GFP*). In the subperineurial glia, septate junctions are formed between two facing subperineurial membranes and appear as a single continuous line along the length of the nerve to create the blood-nerve barrier (**Fig. 6E”**, yellow arrow). With Sdc knock down in the subperineurial glia, both RNAi lines had discrete nerve regions that appeared larger in diameter with a random distribution along each nerve. Within the enlarged regions, the subperineurial glia membranes were expanded, irregular and disorganized (**Fig. 6F-G”,** white arrows), in contrast to the thin and smooth membranes seen in control (**Fig. 6E**’). We observed that septate junction strands formed multiple interconnected strands but these changes were only observed within the enlarged regions (**Fig. 6F”, G”**, yellow arrows) and the septate junctions appeared normal outside these regions. Of interest, the enlargement of the subperineurial glia did not resemble a typical swelling phenotype seen with other mutants such as transporter mutants or ER stress, which cause bulges, fluid accumulation, or visible vacuoles (Leiserson et al., 2011). When quantified (**Fig. 6H**), the prevalence of such an enlargement phenotype was significantly higher with RNAi-1 (*SPG>Dcr2, mCD8::RFP, Sdc RNAi-1*) compared to control (*SPG>Dcr2, mCD8::RFP*) or RNAi-2 (*SPG>Dcr2, mCD8::RFP, Sdc RNAi-2*) which had a small but not significant increase in affected nerves.

We next tested the role of Sdc within the wrapping glia. In controls wrapping glia extend membrane projections wrapping single or bundles of the axons (*Nrv2>Dcr2, mCD8::RFP*) (**Fig. 6I’**, yellow arrow). Sdc knockdown with both RNAi lines resulted in prominent wrapping glia ensheathment defects (*Nrv2>Dcr2, mCD8::RFP, Sdc RNAi-1; Nrv2>Dcr2, mCD8::RFP, Sdc RNAi-2*). Specifically, we observed a loss of the complex glial processes (**Fig. 6J’, K’,** yellow arrows) or breakage of the wrapping glia processes along the length of the nerve. However, the lack of wrapping did not impact axonal morphology (**Fig. 6J”, K”**). With *RNAi-1,* 95.7% of the peripheral nerves showed a lack of ensheathment compared to 46% with *RNAi-2*, and both were significantly higher than control (**Fig. 6L**). Interestingly, the migratory ability of wrapping glia remains intact with Sdc knockdown as the wrapping glia were distributed normally along the nerve (n = 29). Of note, the locomotion defects we observed with Sdc RNAi-1 knockdown in wrapping glia (**Fig. 3**) is likely due to changes in the CNS and not disruption of the peripheral glia, as disruption of wrapping glia ensheathment in the PNS does not alter animal movement significantly (Kottmeier et al., 2020).

To confirm the effects of Sdc knockdown in the wrapping glia, we utilized the heteroallelic combination of *Sdc* mutants and Nrv2 endogenously tagged with GFP (Nrv2::GFP) that specifically labels wrapping glial membranes. We observed normal wrapping glia morphologies in both the heterozygous *Df(2R)48, ubi-Sara/+* (**Fig. 6N**) and control (*w^1118^*) (**Fig. 6M**). In contrast, the heteroallelic combination of *Df(2R)48, ubi-Sara/Sdc^97^* exhibited wrapping glial disruptions similar to those seen with the Sdc RNAi (**Fig. 6O,P**) with 42% of nerves affected, significantly more than control (**Fig. 6P**). Overall, our data suggests that Sdc plays an important role in either the development or maintenance of the wrapping glia ensheathment of peripheral axons.

### Syndecan is required for perineurial glial ensheathment of peripheral nerves

The role of Sdc in the outer layer perineurial glia was the potentially most interesting of all the glial layers given the role of syndecans in mediating cell to ECM interactions. To determine the role of Sdc in perineurial glia, we expressed each Sdc-RNAi along with Dcr2 and a fluorescently tagged cytoskeleton marker (Lifeact::GFP) using the perineurial glial driver, *46F-Gal4*.

Compared to control nerves (*46F>Dcr2, Lifeact::GFP*) (**Fig. 7A**), we detected sections of peripheral nerves with scattered rather than continuous perineurial glial ensheathment and segments of nerve lacking perineurial glial all together and this phenotype was most apparent using RNAi-1 (*46F>Dcr2, Lifeact::GFP*, *Sdc RNAi-1*) (**Fig. 7B-C**, white arrow). On average 97% of RNAi-1 and 55% of RNAi-2 nerves displayed this phenotype, both significantly higher compared to control (4%) (**Fig. 7D**)

**Figure 7:**
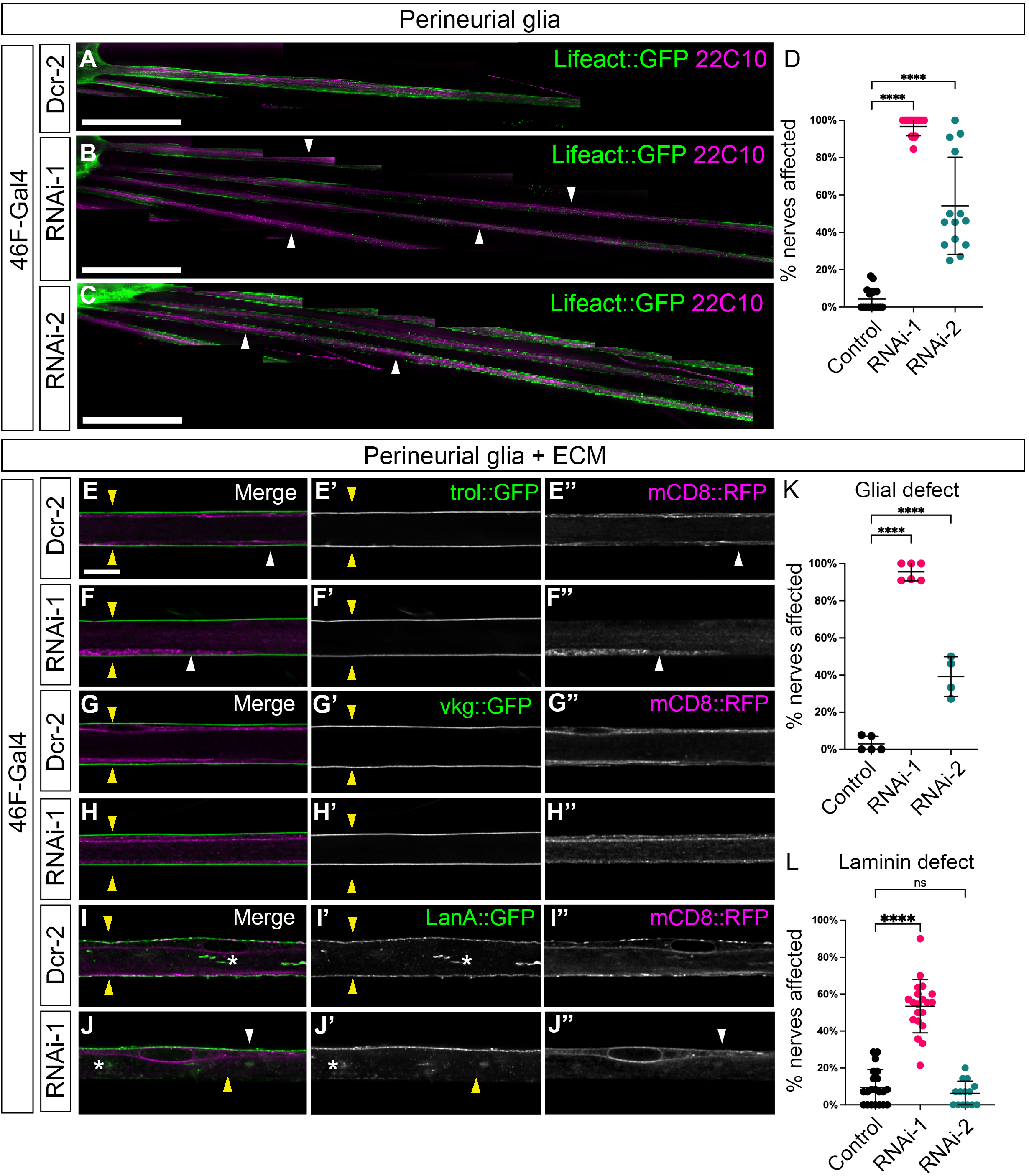
Perineurial glial ensheathment and ECM defects observed with Syndecan knockdown. Loss of Syndecan in the perineurial glial driven using 46F-Gal4. All RNAi lines were co-expressed with Dicer-2. In all graphs, bars indicate the mean ± SD and each represent a One-way ANOVA, Dunnett post-hoc multiple comparison test: **** p<0.0001; ns – not significant. (**A-C**) Representative fluorescent image of the abdominal peripheral nerves at low resolution. *46F-Gal4* was used to express Lifeact::GFP (green) labeling the actin-cytoskeleton network in the perineurial glia and axons were immunolabeled with anti-futsch (magenta). 46F-Gal4 also drove the expression of Dcr2 (**A**), Sdc RNAi-1 (**B**), Sdc RNAi-2 (**C**). Note the segments of the peripheral nerve lacking coverage of perineurial glia (white arrows) in both RNAi knockdown groups (**B-C**). Scale bar, 100 µm. (**D**) Quantification of perineurial glia disruption. A defective nerve was defined as a segment of the nerve within the nerve extension region showing glial distributions to only one side of the nerve or the absence of glia. Both RNAi lines (Sdc RNAi-1, Sdc RNAi-2) have significantly increased perineurial glia defects compared to control (Dcr2). (**E-K**) Representative longitudinal section of peripheral nerve labeled for the ECM proteins Viking (**E-F**), Perlecan (**G-H**) and Laminin-A (**I-J**), all endogenously tagged GFP (green). Membrane-bound RFP (mCD8::RFP, magenta) was driven using 46F-Gal4 to visualize perineurial glia membranes. Loss of Sdc led to glial ensheathment defects (**F”, J”,** white arrows) where the glial membrane is found on only one side of the nerve compared to control where the glial surrounds the entire nerve (**E”,** white arrows). Loss of Sdc also lead to a reduction in the levels of Laminin-A (**J’**, yellow arrows) compared to the uniform distribution in controls (**I’**, yellow arrow). The interior deposition of Laminin-A was not disrupted (**I’, J’**, asterisks). Viking and Perlecan had normal distributions in both control and RNAi lines (**F’, H’**, yellow arrows). Scale bar, 10 µm (**K**) Quantification of the ensheathment defects of perineurial glia. Loss of Sdc with either RNAi line (Sdc RNAi-1; Sdc RNAi-2) led to a significant increase in the number of nerves with loss of complete nerve ensheathment compared to control (Dcr2). (**L**) Quantification of the loss of Laminin-A expression. Sdc RNAi-1 had a significant increase in the number of nerves with loss of Laminin-A expression on one or both sides of the nerve compared to control (Dcr2) and Sdc RNAi-2.

We next assessed the effects on the formation of the glial sheath using the membrane mCD8::RFP marker in the context of the overlying ECM marked with Perlecan, Collagen or Laminin (**Fig. 7D-I**). We found that with both RNAi lines (*46F>Dcr2, mCD8::RFP, Sdc RNAi-1; 46F>Dcr2, mCD8::RFP, Sdc RNAi-2*) the perineurial glia failed to wrap around the peripheral nerve with processes remaining limited to one side of the nerve (**Fig. 7F”, J”**; white arrows) compared to encompassing the entire nerve in the control (*46F>Dcr2, mCD8::RFP)* (**Fig. 7E**, white arrows). When quantified both RNAi lines had significantly greater numbers of nerves affected per animal (on average: RNAi-1 – 96%; RNAi-2 – 39%) compared to control (**Fig. 7K**). Therefore, Sdc plays a critical role in the migration of perineurial glia and the creation of the full glial wrap around each peripheral nerve.

*In vitro*, syndecan deficiency is often observed with disorganized ECM fibers (Klass et al., 2000; Yang and Friedl, 2016). With the reduction in the perineurial glia, we were curious to investigate the impact loss of Sdc would have on the ECM. We focused on three common ECM components that were previously identified and analyzed in the *Drosophila* neural lamella (Xie and Auld, 2011; Clayworth and Auld, 2025), perlecan (Trol), collagen-IV (Viking), and laminin (LanA), all endogenously tagged with GFP. Normally, the ECM appears as a continuous uniform sheath surrounding the surface of the nervous system as demonstrated by a single line of Perlecan and Collagen in nerve longitudinal sections (**Fig. 7E’, G’, I’,** yellow arrows). Perlecan and Collagen distribution within the nervous system was not different between control and Sdc knockdowns. Continuous, uninterrupted ensheathment was observed in all groups (**Fig. 7F’, H’,** yellow arrows), despite the presence of the perineurial ensheathment defects. We next assayed laminin deposition using a GFP tagged laminin alpha subunit (LanA::GFP). Laminin is deposited by both the perineurial and wrapping glia and normally forms a uniform sheath around the exterior of the nervous system and with concentrations within the center of the peripheral nerve (Petley-Regan et al, 2016; Clayworth and Auld, 2025) (**Fig. 7I**, yellow arrows). With Sdc *RNAi-1* within the perineurial glia (*46F>Dcr2, mCD8::RFP; Sdc RNAi-1*), we detected segments of the nerve that had reduced or lacked laminin distribution along the exterior of the nerve (**Fig. 7J’**, yellow arrow). With RNAi-2 *(46F>Dcr2, mCD8::RFP; Sdc RNAi-2)* laminin appeared unaffected. When quantified, only RNAi-1 showed a significant percentage of nerves with laminin defects (53% on average) compared to control (*46F >Dcr2, mCD8::RFP*) (**Fig. 7L**). Thus, the loss of Sdc affects perineurial glial distribution and ensheathment along the peripheral nerve as well as the glial deposition of laminin in the neural lamella surrounding the nerve.

### Loss of Syndecan reduces the number of peripheral perineurial glia

The loss of the perineurial sheath with Sdc knockdown could be due either to an inability to extend sufficient membrane to wrap around the nerve or to a reduction in the number of perineurial glia. To test whether loss of Sdc affects the number of perineurial glia, we quantified the number of perineurial nuclei on the A8 (8^th^ abdominal) nerve using GFP tagged with a nuclear localization signal (NLS::GFP) driven by *46F-Gal4*. There are on average 40-50 perineurial glial evenly distributed along the A8 nerve in the third instar larvae. With Sdc knockdown using RNAi-1 (*46F>Dcr2, NLS::GFP*, *Sdc RNAi-1*), we observed the perineurial glia population reduced to 40% of control (*46F>Dcr2, NLS::GFP*) (**Fig. 8A-D**). Oddly, we found a small but significant increase in the perineurial glia population with RNAi-2 (*46F>Dcr2, NLS::GFP, Sdc RNAi-2)* to 113% of control (**Fig. 8D**). Thus, the lack of defects in laminin deposition with RNAi-2 compared to RNAi-1 could be a function of the reduced glial presence with *RNAi-1* compared to *RNAi-2,* which has a slight increase in glial cells. This increase in perineurial glia might also reflect the increase in locomotion parameters we observed with PG knockdown using RNAi-2 (**Fig. 3**).

**Figure 8:**
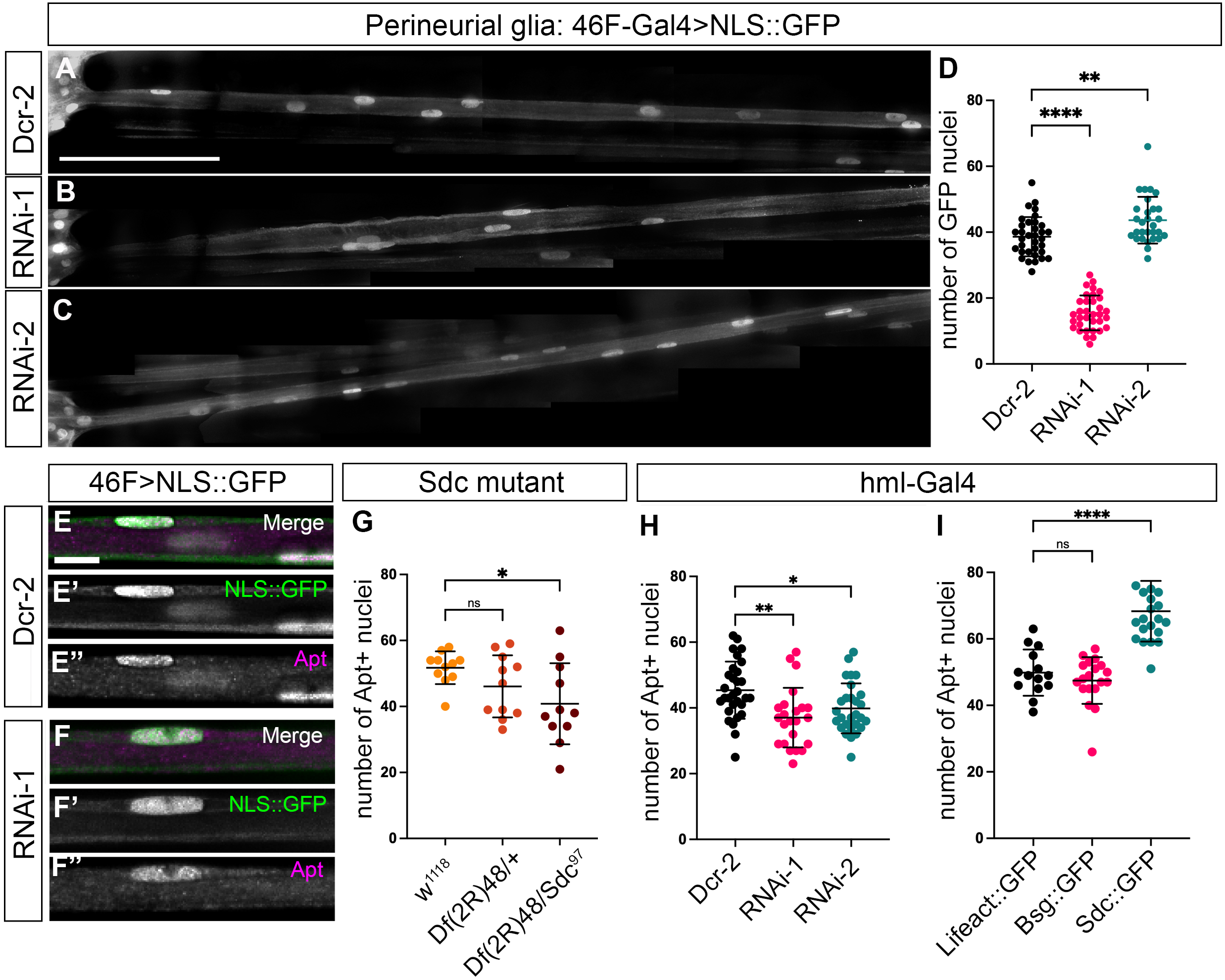
Syndecan expression levels in both hemocytes and perineurial glia affect perineurial glia numbers in the PNS. Analysis of the cell autonomous (A-D) and non-autonomous (H-1) effects of Sdc expression changes. All RNAi lines were expressed along with Dicer2. Nuclei were counted along the abdominal 8 (A8) nerve. On all graphs, bars indicate the mean ± SD, with the mean nuclei count analyzed using a One-way ANOVA, Dunnett post-hoc multiple comparison test: **** p<0.0001; **p<0.01; *p<0.05; ns – not significant. (**A-C**) Representative cross-section images of peripheral nerves with 46F-Gal4 driving the expression of nuclear localized GFP (NLS::GFP) in the presence of Dcr2 (A), Sdc-RNAi (B) and Sdc RNAi-2 (C). Both RNAi lines were co-expressed with Dicer-2. Scale bar, 100 μm (**D**) Quantification of GFP+ nuclei count. Sdc RNAi-1 had significantly less while Sdc RNAi-2 had more nuclei compared to control (Dcr2). (**E-F**) 46F-Gal4 driven expression of NLS::GFP (green) and immunolabeling for Apontic (magenta) were similar between control (Dcr2) (**E**) and Sdc RNAi-1+Dcr2 (**F**). Scale bar, 10 um (**G**) Quantification of the number of Apontic positive nuclei along the A8 nerve in *Sdc* homozygous mutants. *Df(2R)48, ubi-Sara/Sdc^97^* had a significant reduction in the number of Apontic positive nuclei when compared to control (*w^1118^*). *Sdc* heterozygous mutant (*Df(2R)48, ubi-Sara/+*) did not. (**H**) The hemocyte driver (hml-Gal4) was used to drive expression of Sdc RNAi and both RNAi-1 and RNAi-2 had a small but significant decrease in Apontic nuclei along the A8 nerve. (**I**) The hemocyte drier (hml-Gal4) was used to drive Lifeact::GFP, Basigin::GFP or Sdc::GFP. Expression of Sdc::GFP lead to a significant increase in the number of Apontic nuclei compared to controls.

The increase in PG numbers with the weaker RNAi-2 line suggested a possibility that the levels of Sdc within the peripheral glia may mediate differential functions. To test the effect of a global loss of Sdc, we utilized the *Sdc* loss of function alleles and tested for the presence of the perineurial glia using apontic immunolabeling. Apontic is a transcription factor that specifically labels the perineurial glia in the peripheral nerve (**Fig. 8E**) (Zülbahar et al., 2018). We found that Apontic expression is not affected by the knockdown of Sdc (**Fig. 8F**). Similar to the knockdown of Sdc using RNAi-1, the heteroallelic combination (*Df(2R)48, Ubi-Sara/Sdc^97^*) had significantly less Apontic positive nuclei (80%) compared to the deficiency alone (*Df(2R)48, Ubi-Sara*/+) or the control (*w^1118^*) (**Fig. 8G**). Thus, systemic loss and a strong reduction in Sdc leads to a reduction in the number of perineurial glia while a small degree of knockdown within the perineurial glia knockdown had the opposite effect.

This leads to a model whereby the expression level of Sdc within the glial layer could be informative to perineurial glia numbers and development and suggests a potential non-autonomous role for Sdc in the perineurial glia. The extracellular domain of Syndecan (ectodomain) can be released by proteolytic cleavage, yielding a soluble proteoglycan that retains binding properties and receptor specificity. For instance, ectodomains of mammalian Syndecans and *Drosophila* Syndecan are constitutively shed from cultured cells (Kim et al. 1994; Spring et al, 1994). Thus, we wanted to test the effect the loss of Sdc in other tissues would have on the perineurial glia. Multiple basement membrane components including proteoglycans are secreted by hemocytes in Drosophila and this process in embryos is essential for the development of the basement membranes (Matsubayashi et al. 2017). One external source of Sdc could be from circulating hemocytes (blood cells). We expressed both RNAi lines within hemocytes (*hml>Dcr2, Sdc-RNAi-1*; *hml>Dcr2, Sdc-RNAi-2*) and observed that loss of Sdc from the hemocytes lead to a small but significant decrease in perineurial glia with RNAi-1 and RNAi-2 (on average 81% and 88% respectively) (**Fig. 8H**). These decreases were not as strong as observed with the PG driver suggesting a mild non-autonomous effect. To further test the non-autonomous effect of Sdc on PG, we overexpressed Sdc within the hemocytes using an unrelated transmembrane protein Basigin as a control (*hml>Bsg::GFP*) or an actin control (*hml>LifeAct::GFP*). The overexpression of Sdc (*hml>Sdc::GFP*), but neither of the controls, led to a significant increase in the number of apontic positive perineurial glia (on average 137% of control) (**Fig. 8I**). Our results suggest that ectopic Sdc either directly or indirectly leads to increased perineurial glial numbers and that Sdc levels play important roles in perineurial glial function and development.

### Syndecan and integrins indirectly interact to mediate perineurial glia ensheathment

The loss of Sdc appeared to have to differential effects on the perineurial glia: a reduction in cell number and a lack of ensheathment of the axon. These phenotypes were highly reminiscent of the effects of due to loss of integrin and focal adhesion complex components. Specifically, RNAi to the beta subunit of the integrin heterodimer (*myospheroid*) and *mys* mutants disrupt the perineurial glial sheath and reduce cell numbers (Xie and Auld, 2011). Focal adhesion formation during cell spreading and migration depends on the cooperation of integrin and cell-surface proteoglycans. For example, vertebrate α_2_β_1_and α_6_β_4_integrins cooperate with syndecans during adhesion to laminin (Hozumi et al., 2006; Ogawa et al., 2007). Given the widespread expression of Sdc at the glial-ECM interface, we tested whether Sdc co-localizes with integrins within the peripheral nerve. We analyzed the distribution of Sdc::GFP compared to the integrin beta subunit Myospheroid (aka βPS). Within the peripheral nerve, Sdc puncta did not co-localize with the integrin positive belt-like regions (**Fig. 9A**, white arrows) and when quantified the Pearson’s coefficient was on average 0.26 ± 0.03 (n = 4 larvae) suggesting that Sdc and integrins do not associate at focal adhesions in Drosophila peripheral glia.

**Figure 9:**
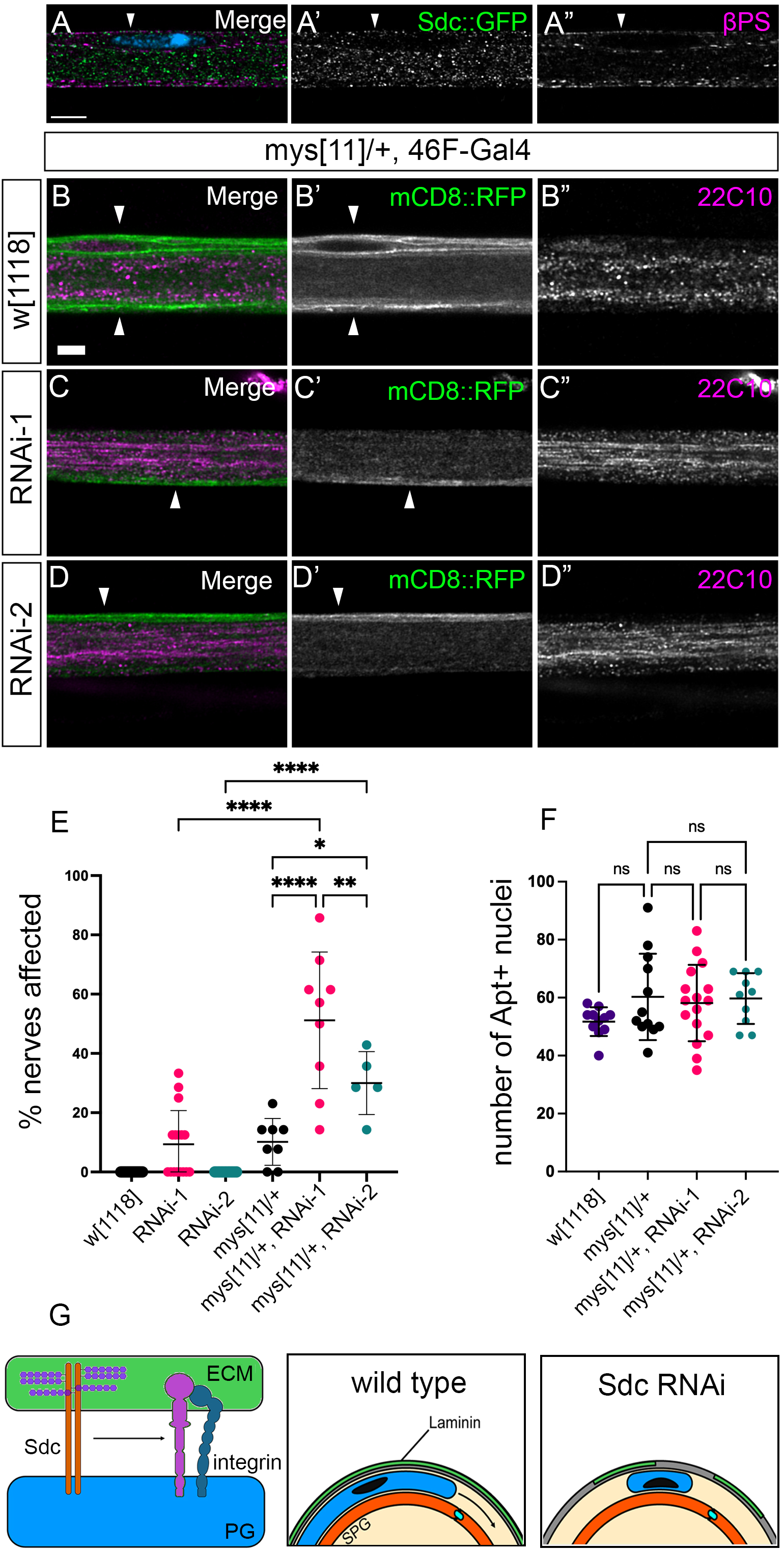
Syndecan and beta-integrin genetically interact to mediate perineurial glial ensheathment but not numbers. (**A**) Representative cross-section images of peripheral nerves Sdc::GFP (green) and immunolabelling for the integrin β-subunit, Myospheroid (βPS, magenta). Sdc and Mysospheroid do not colocalize (white arrow). Scale bar, 10 μm (**B-F**) Sdc-RNAi mediated knockdown (in the absence of Dicer2) paired with a heterozygous loss of function *mysopheroid* mutation (*mys^11^*). (**B-D**) Representative cross-section images of peripheral nerves with 46F-Gal4 driving mCD8::RFP (green) to label glial membranes and immunolabelling for futsch (22C10, magenta). Perineurial glial in both Sdc RNAi-1 and RNAi-2 in combination with *mys^11^* failed to ensheath the entire peripheral nerve (**C’, D’**, white arrows) compared to control (*w^1118^*) (**A’**, white arrows). Scale bar, 10 μm (**E**) Quantification of glial ensheathment defects. Sdc RNAi-1 and RNAi-2 in combination with *mys^11^*/+ had significantly more glial ensheathment defects compared to each RNAi line alone (in the absence of Dcr2), *mys^11^/+* alone or control (*w^1118^*). Note: only significant differences are displayed; non-significant differences are not. (**F**) Quantification of the number of Apontic nuclei on abdominal A8 nerve. There were no differences between Sdc RNAi-1 and RNAi-2 in combination with *mys^11^*/*+* compared to *mys^11^/+* alone or control (*w^1118^*). On all graphs, bars indicate the mean ± SD, analyzed using a One-way ANOVA, Dunnett post-hoc multiple comparison test: **** p<0.0001; *** p<0.001; ** p<0.01; *p<0.05; ns – not significant. (**G**) Model of Syndecan function in perineurial glia. Syndecan interacts in an indirect manner with the focal adhesion complex (integrins: alpha and beta subunits) in the membrane of the perineurial glia. This interaction may regulate the association of focal adhesions with the overlying ECM to direct PG ensheathment in wild type. Loss of Sdc leads to reduced adhesion and loss of the perineurial glial sheath.

While Drosophila Sdc did not localize with focal adhesions in the peripheral nerve, we wanted to test if there was a genetic interaction. We tested the ability of a loss of function allele of the beta-integrin subunit (*mys^11^*) to enhance the Sdc RNAi knockdown. In the absence of Dcr2, Sdc RNAi-1 had a reduced penetrance of the ensheathment phenotypes while Sdc RNAi-2 had none (*46F>mCD8::RFP, Sdc RNAi-1; 46F>mCD8::RFP, Sdc RNAi-2*) (**Fig. 9E**). However, when placed in a *mys^11^/+* heterozygous background, ensheathment defects were observed where the glial fail to cover the full circumference of the peripheral nerve (**Fig. 9C’, D’**, white arrowheads). When quantified (**Fig. 9E**), there was a significant increase in the degree of ensheathment defects with both RNAi lines (*mys^11^/+, 46F>mCD8::RFP, Sdc RNAi-1; mys[11]/+, 46F>mCD8::RFP, Sdc RNAi-*2) compared to the controls (*mys^11^/+,46F>mCD8::RFP; 46F>mCD8::RFP, Sdc RNAi-1; 46F>mCD8::RFP, Sdc RNAi-*2).

Of note there were no significant changes to the number of perineurial nuclei (**Fig. 9F**) suggesting that interactions between integrin and Sdc affect the production of the glial wrap around the peripheral nerve but not proliferation. These results suggest two roles for Sdc in perineurial glia – regulation of the glial sheath around each peripheral nerve likely in concert with focal adhesions (**Fig. 9G**) and regulation of perineurial glial numbers through an unknown mechanism.

## Discussion

Syndecan, a transmembrane heparan-sulfate proteoglycan, has been postulated as a key player in cellular communication, yet it’s *in vivo* impact in nervous system function specifically with respect to glial function remains largely untested. We have delineated two aspects of Sdc function in glia: Sdc regulates neuroblast proliferation; Sdc regulates the ensheathment processes of multiple glial types. Overall, this suggests Sdc acts as a fundamental regulator of key cellular processes in *Drosophila* glia.

### Syndecan is required in CNS glia

We observed a range of CNS defects including elongation and a loss of brain lobe size. The elongation of VNC seen with pan-glial knockdown of Sdc indicates a possible contribution to cell-ECM adhesion. Elongation of the VNC is often observed with disruption in glial-ECM contact (Meyer et al., 2014; Skeath et al., 2017). Loss of integrin in the perineurial glia leads to dramatic elongation of the VNC (Xie and Auld, 2011; Meyer et al., 2014) far beyond that observed with the Sdc knockdown down. This suggests Sdc likely facilitates integrin mediated glia-ECM adhesion in the CNS but is not the key mediator. The CNS elongation was only observed with RNAi-mediated knockdown and not in the systemic loss in the *Sdc* mutants. We are confident that the Sdc RNAi lines are specific and this is not due to an off-target effect. Rather this observation may result from the differential expression of Sdc in the glial layers of the CNS rather than the global loss.

The reduction in brain lobe size points to a potential role for glial Syndecan in regulating the neuroblasts. In other systems, Syndecan plays an important role in stem cell survival and stem cell properties. In Drosophila, in the midgut loss of Sdc alters stem cell and nuclear shape and leads to loss from the tissue, while loss of Sdc in neural stem cells generates nuclear envelope defects during cell division (Eldridge-Thomas et al., 2025). Of note, Sdc-RNAi knockdown in neuroblasts did not affect brain size (Eldridge-Thomas et al., 2025), while in this study loss of Sdc in all glial significantly reduced brain lobe diameter due to a reduction in neuroblast proliferation. Given that the trans-heterozygous Sdc mutants also had reductions in brain lobe size, this suggests that the glial Sdc contribution is the major reason for this defect. The reduced neurogenesis observed in pan glial Sdc knockdown may have several causes. One possibility could be that Sdc functions in the cortex glia or the subperineurial glia that are associated with the neuroepithelium. A sub-population of cortex glia directly contact the neuroblasts, transition zone, and neuroepithelium of the optic lobe and regulate the proneural wave progression that drives the neuroepithelium to neuroblast transition (Morante et al., 2013). The cortex glia can regulate the neuroepithelium through trophic factors, such as the EGFR ligand, Spitz, which triggers the neuroblast transition (Morante et al., 2013). As well cortex glia respond to other growth factors that promote growth and proliferation, including FGF, IGF, and PDGF (Avet-Rochex et al., 2012, 2014; Read, 2018). Syndecans are necessary in the activation of many trophic factor signaling cascades by regulating the availability of growth factors including EGF, FGF and VEGF (Afratis et al., 2017; Corti et al., 2019; Kwon et al., 2012; Mahtouk et al., 2006; Wang et al., 2015). Cortex glia also express Dilp-6 to regulate neuro-proliferation through the establishment of a stem cell niche (Read, 2018; Harrison et al., 2021). Sdc can directly affect IGF signaling as a *Sdc* hypomorphic mutation have lower expression of Dilp-2 in the brain (De Luca et al., 2010). Similarly, vertebrate syndecan has been implicated in IGF signaling by mediating crosstalk between insulin-like growth factor-1 receptors and integrin complexes to direct cell migration and invasion (Beauvais and Rapraeger, 2010; Rapraeger et al., 2013). Thus, Sdc could indirectly influence neuroblast proliferation in the brain lobe by controlling cortex glial proliferation or conversely changes to glial Sdc could directly affect the disruption of growth factors available for neuroblast proliferation.

### Syndecan is required through all three peripheral glial layers

Syndecan has a clear role in the peripheral nervous system such that downregulation of Sdc in all three layers lead to a disruption in glial morphology. Both the SPG and WG had normal cell numbers and glia were localized to their normal locations with the nerve suggesting a role for Sdc in sheath formation or maintenance in these layers. Of interest that Sdc likely has two roles in the perineurial glia, proliferation and migration/sheath formation.

Perineurial glia are born at the CNS/PNS transition zone in the first larval instar stage and proceed to proliferate and then migrate into the periphery (von Hilchen et al, 2013). Migration is followed at later stages by the spread of the glial membranes to surround each peripheral nerve a process that is normally complete by the 3^rd^ instar stage. Therefore, we propose that Sdc has different temporal distinct functions in the perineurial glia: proliferation and migration leading to ensheathment. One possibility is that the absence of Sdc prevents perineurial glia from interacting or binding effectively with the overlying ECM. We observed breakages in laminin deposition upon knockdown of Sdc in the perineurial glia of the PNS. Perineurial glia secrete and can potentially deposit laminin (specifically the laminin LamaninA:LanB1:LanB2 complex) into the surrounding neural lamella (Petley-Ragan et al., 2016). In other cell types, syndecan knockdown does not impact laminin production and secretion, but rather disrupts the extracellular assembly and the arrangement of the ECM architecture for the ECM-protein producing cells (Klass et al., 2000; Yang and Friedl, 2016). Drosophila Syndecan binds to LamininA (LanA) (Ryo et al., 2004). This leads to the suggestion that the absence of Sdc either: 1. disrupts perineurial glia numbers/migration and thus the deposition of laminin, or 2. disrupts the assembly and arrangement of laminin the ECM, which affects the perineurial glia and sheath formation.

Syndecan is also important to regulate the number of perineurial glia, a result observed with both the strong RNAi and the *Sdc* mutants. An unexpected observation was the differential effects of Sdc RNAi-1 and RNAi-2 on the number of perineurial glia. Similar to our observations in the CNS, this could be due to differences in the Sdc knockdown in terms of the remaining Sdc revealing different functions within the peripheral nerve. We don’t think that off-target effects cause this differential result as both RNAi lines have been extensively used and verified. Rather the HS-GAG chains of syndecan can interact with ECM proteins such as laminin or fibronectin to sequester/enhance or quench interactions (Hoffman et al., 1998; Woods et al., 2000). This later point might explain the differential effects that we observed between strong and weak Sdc knockdown where strong loss of Sdc (RNAi-1 or *Sdc* mutants) resulted in reduced PG numbers while a weak knockdown increased PG numbers. Strong loss of function may lead a lack of ligands or signaling to promote glial proliferation, while weak loss of function may directly open up potential ligands due to a reduction in the quenching of ECM interactions to allow for more signaling. The differential functions of Sdc might also explain the effect of overexpression of Sdc within the blood cells (hemocytes). The ectodomain of Sdc can be shed from originating cells (Kim et al. 1994; Spring et al, 1994) and soluble syndecan ectodomains can compete with intact syndecans for extracellular ligands (Steinfeld et al. 1996). Loss of Sdc from hemocytes lead only to a mild reduction in PG numbers suggesting that lower circulating levels of Sdc did not have a strong effect. Conversely increasing Sdc expression in hemocytes dramatically increased PG numbers suggesting either increased Sdc shedding affected perineurial glia division directly or indirectly by altering hemocytes themselves to indirectly affect the perineurial glia. The identity of the mechanism control PG number remains to be discovered.

Our experiments found loss of Sdc leads to ensheathment defects in perineurial glia. In the context of perineurial glia, Sdc is highly expressed and localized to glial-ECM boundaries suggesting a role for Sdc as a mediator of cell-ECM interaction. In the CNS, the extension of the VNC length in the absence of Sdc matches observed phenotypes seen with loss of integrins and degradation of the ECM albeit to a lesser extent (Meyer et al., 2014; Skeath et al., 2017). In the PNS, we observed the ensheathment of the perineurial glia was disrupted in that each perineurial glia was limited to one side of the nerve or absent and this was observed in both weak and strong RNAi lines. These results were similar to those observed with the loss of focal adhesion components (the β-subunit of integrin and Talin) or after degradation of the ECM (Xie and Auld, 2011). Syndecan has been demonstrated to provide a mechanical linkage from the ECM to the cytoskeleton through binding to laminin via the HS-GAG chains and the actin-network through Ezrin or α-actinin (Granes et al., 2000; Yamashita et al., 2004; Carulli et al., 2012; Okina et al., 2012). Syndecan can also act synergistically with integrin to regulate ECM adhesion, focal adhesion assembly, and cytoskeletal re-arrangement (Saoncella et al., 1999; Morgan et al., 2007; Okina et al., 2012; Fiore et al., 2014). Vertebrate Syndecan-4 has been identified as a cellular tension sensor whereby application of external tension on syndecan-4 drives integrin-based focal adhesion growth (Chronopoulos et al., 2020).

We do not think that Sdc functions in the same protein complex as integrin. Using super-resolution imaging, we observed that Sdc does not co-localize with integrin in the peripheral nerves of *Drosophila*. This is in contrast to reports of syndecan-4 and integrin, having direct interactions via the syndecan ectodomain and cytosolic domain (Wang et al., 2010). The spatial organization of syndecan and integrin could reflect a model of syndecan recruitment postulated by Roper et al. (2012). In this model, Syndecan is present at the initial stages of integrin focal adhesion complex formation and subsequently syndecan is repelled from the focal adhesion as the focal adhesion matures. Alternatively, the long GAG chains on syndecan could influence integrin complex association with the ECM even if the protein cores do not interact (Roper et al., 2012; Stepp et al., 2015). Given that the loss of the beta subunit enhanced both Sdc RNAi lines without the presence of Dicer2, this suggests that there is synergistic interaction between focal adhesions and Sdc in PG glia but the mechanism that underlies this interaction remains to be discovered.

## Summary

In conclusion, we have characterized two potential glial functions for Sdc *in vivo* during *Drosophila* neural development. Knockdown of Sdc reduces neuroblast proliferation within the optic lobe causing a severe reduction in brain lobe size. We also found that loss of Sdc is associated with a disrupted distribution of perineurial glial processes along the length of the nerve, in combination with changes in the deposition of laminin. Our findings provide the first evidence for Sdc function in PNS glia, along with its potential implication in neural stem cell maintenance and emphasizes the advantages of using of *Drosophila* as a platform to study Sdc *in vivo* function, as there are no functional redundancy challenges to investigating as there are with vertebrates.

## Material and Methods

### Fly strains and genetics

Standard *Drosophila* husbandry techniques and genetic methodologies were used to obtain files of the required genotypes for each experiment from the original transgenes. For the original fly strains used in this study see **Table 1**. All crosses were carried out 25°C with Dcr-2 in the background unless indicated otherwise.

**Table 1:**
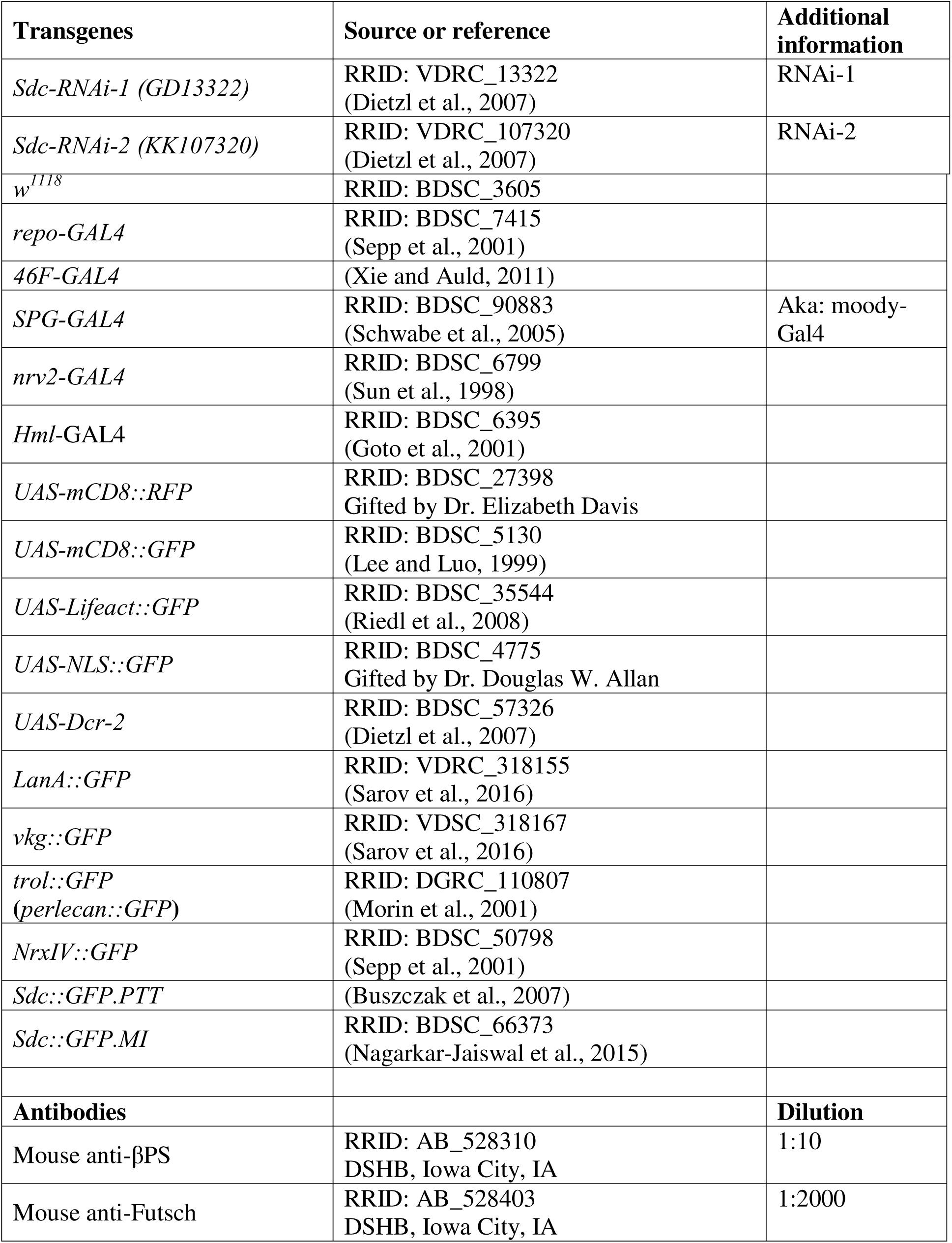
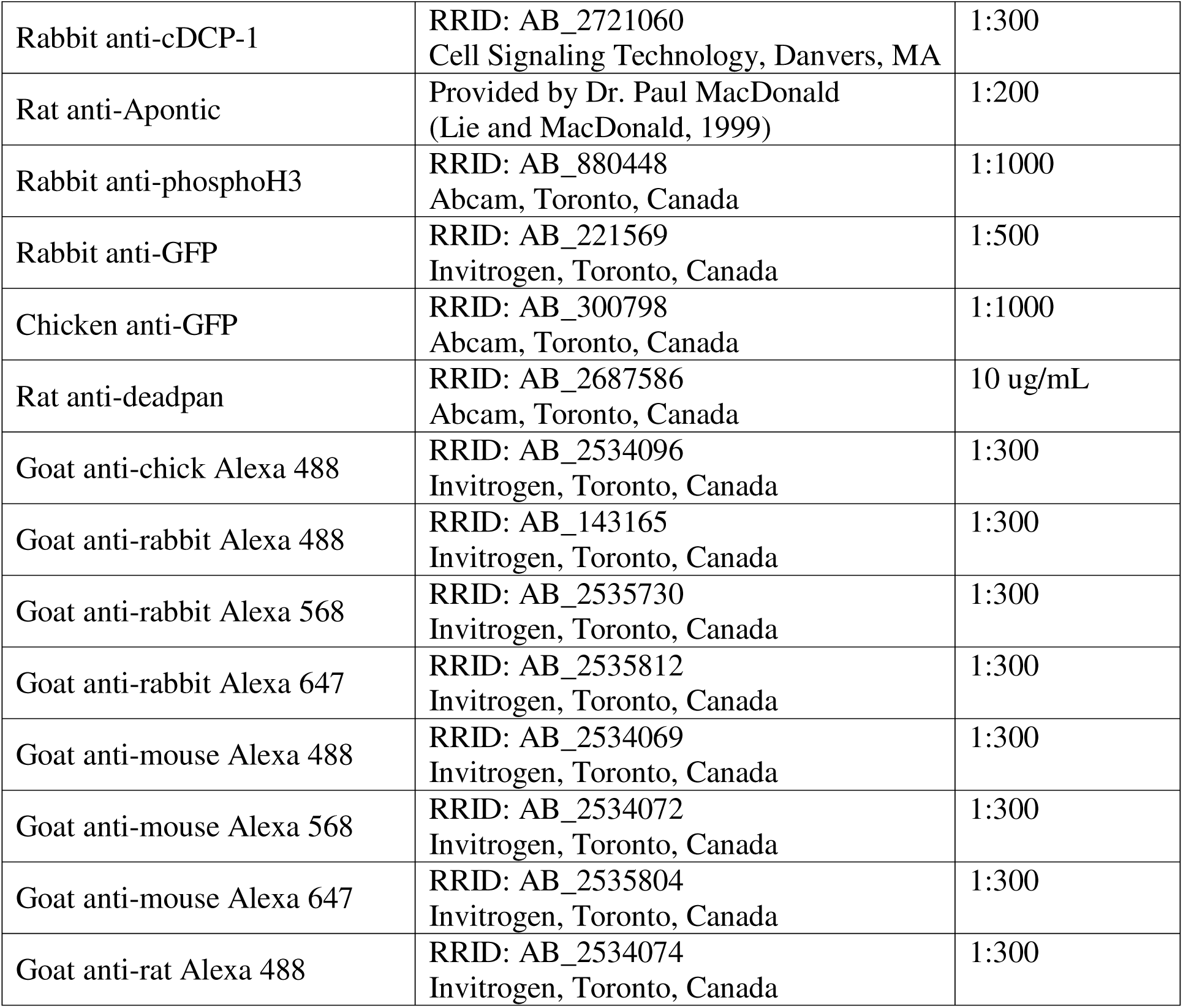
Origins of the transgene and antibodies used in this study.

### Larvae dissection and immunofluorescence

Third instar larvae were filleted and dissected in PBS on Sylgard plates. The protocol for fixation of larvae was in 4% paraformaldehyde with 1X PBS for 20 minutes. For integrin immunolabelling, larvae were fixed for only 10 minutes. Fixed fillets were washed two times in 1X PBS for 5 minutes each then washed three times in PBST (1X PBS, 0.1% Triton X-100) for 10 minutes each at room temperature. For immunofluorescence, samples were first transferred into blocking solution (4% heat-inactivated goat serum, 0.1% PBST) at 4°C overnight. All antibodies were diluted in blocking solution. For primary antibody incubation, samples were placed on an orbit shaker at 4°C overnight. Larvae fillets were then washed three times with PBST for 10 mins each and incubated with secondary antibody solution for two hours at room temperature. Lastly, samples were washed with PBST three times for 10 minutes each. Samples were equilibrated via glycerol series up to 90% glycerol and mounted with Vectashield antifade mounting medium (Vector Laboratories, Burlington, Canada). DAPI (1:1000 of 1 ug/mL) (Thermo Scientific) was added along with the secondary antibodies.

### Imaging and Image processing

Fluorescent images of the peripheral nerves were taken with a DeltaVision microscope (Applied Precision, Mississauga, Canada) using a PlanApoN 60X oil immersion objective (NA=1.42) at 0.2 µm steps in the z-direction. Individual channels were deconvolved separately and merged back together with SoftWorx image processing software (SoftWorx, Toronto, Canada). Deconvolution is performed using a point spread function measured with a 0.2 µm fluorescent bead (Invitrogen, Toronto, Canada) in Vectashield Mounting Medium. The number of deconvolution iterations was set to terminate once the normalized error criterion for the constrained iterative algorithm begins to plateau. Orthogonal sections were generated using SoftWorx. Images of peripheral nerves were stitched using the Pairwise Stitching plugin (Preibisch et al., 2009) in Fiji (Schindelin et al., 2012). Images were further processed and compiled using Adobe Photoshop and Adobe Illustrator (Adobe Creative Cloud).

To measure the ventral nerve cord to body length ratio, ventral nerve cords were imaged using a DeltaVision microscope (Applied Precision, Mississauga, Canada) with a Plan 20X air objective (NA=0.4) at 1 µm steps in the z-direction. Body length was captured using an Axioplan-2 microscope (Carl Zeiss Canada, Toronto, Canada) with a Zeiss A-Plan 2.5X objective (NA=0.06) air objective. Images of VNC and larval body were stitched using the Pairwise Stitching plugin (Preibisch et al., 2009) in Fiji. Images were further processed and compiled using Adobe Photoshop and Adobe Illustrator (Adobe Creative Cloud).

To measure the effects of Sdc loss in the brain lobes, fluorescent images of brain lobes were acquired with an Olympus FV1000 Laser Scanning Confocal Microscope (Bioimaging Facility, UBC, Vancouver, Canada) using a UPLSAPO 30X silicone oil immersion objective (NA=1.05) at 0.73 µm steps. Digital zoom was optimized to ensure Nyquist sampling. Image stitching was achieved via the Pairwise Stitching plug-in (Preibisch et al., 2009) in Fiji (Schindelin et al., 2012). Images were further processed and compiled using Adobe Photoshop and Adobe Illustrator (Adobe Creative Cloud). To estimate neuroepithelial volume for each Z slice the outer boundary of the neuroepithelium was manually traced using immunolabeling for β-catenin (Arm). The area was measured for each slice across the entire Z-stack to calculate the neuroepithelial volume. In this volume the number of mitotic nuclei were counted using immunolabeling for phosphoHistone3 channel. The proliferation extent was calculated by the number of mitotic cells within the calculated neuroepithelial volume.

Super-resolution images were acquired with a Zeiss LSM800 Confocal with Airyscan (Carl Zeiss Canada, Toronto, Canada) using a Plan-APOCHROMAT 63X oil immersion objective (NA = 1.4) at 3.5X digital zoom. Z-step collection and frame size was set to ensure Nyquist sampling. Laser intensity and master gain of each channel were optimized to ensure intensity spans 1/3 – 2/3 of the displayed histogram. Scan speed was set to 4 –7 µs/pixel. Raw images were further processed using ZEN 3.1 (blue edition, Carl Zeiss Canada, Toronto, Canada) and compiled using Adobe Photoshop and Adobe Illustrator (Adobe Creative Cloud).

### Larval tracking

Larval tracking was carried out as previously described (Das et al., 2023). For each larval tracking session, multiple 3^rd^ instar larvae were added to a fresh 2% agar plate. Food-safe navy-blue dye was added to enhance contrast between larvae and agar. Larval movements were recorded continuously for 60 seconds using a Canon VIXIA HF R800 video camera (Canon). The recorded movies were analyzed using Fiji plug-in wrmTrck (Nussbaum-Krammer et al., 2015; Brooks et al., 2016) to calculate speed and travel distance.

### Larval survival assay

A set number of third instar larvae of the desired genotype were placed into fresh food vials. The number of pupae within each vial was counted after larvae ceased wandering. Each vial was then observed daily for two weeks after the first adult eclosed. The newly eclosed adults were separated by sex and observed for two more days. Adults that died within this 48-hour window were considered to be adult lethal.

### Statistical analyses

All statistical analyses were conducted using GraphPad Prism 9 (GraphPad Software, La Jolla, CA). For larval locomotion assay (scatter plots) each point on the graph represents a single 3^rd^ instar larvae. For all peripheral glial morphological analyses, 6-8 nerves were assessed per larvae and each point on the graph represents the percentage of nerves affected in an individual animal. For all peripheral glial nuclei analyses, the number of nuclei (GFP or Apontic labelled) were counted on the abdominal 8^th^ nerve (A8) and each point on the graph represents the number of nuclei per nerve. For all graphs, the difference in means of control and experimental were analyzed using an ordinary parametric one-way ANOVA with a Tukey *post hoc* test for comparison to the control or a Dunnet’s post host test for comparison across all genotypes. All graphs show mean and standard deviation (SD).

## Acknowledgements

We would like to thank Dr. Mary Gilbert for constructive comments on this manuscript. We thank Dr. Paul MacDonald for the anti-apontic antibody and Dr. Matt Ramer for use of the Airyscan confocal. We want to thank the core imaging facilities in core facilities provided by the Life Sciences Imaging Facility and the Biosciences Imaging Facility. Fly stocks were obtained from the Bloomington Drosophila Stock Center (NIH P40OD018537) and from the Vienna Drosophila Resource Center. Monoclonal antibodies were obtained from the Developmental Studies Hybridoma Bank, created by the NICHD of the NIH and maintained at The University of Iowa, Department of Biology, Iowa City, IA 52242.

