## Supplementary Figure 1 for "Syndecan is critical for *Drosophila* CNS and PNS glia function"

**Supplemental Figure 1: Sdc RNAi leads to loss of Sdc::GFP in CNS and PNS glia**

**(A-C)** Knockdown of Sdc using the pan-glial repo-Gal4 driver with glial membranes marked with mCD8::RFP (magenta) and Sdc tagged with GFP (Sdc::GFP, green). Scale bars: 20 µm A,C; 10 µm B,D.

(**A,B**) Control brain lobe (**A**) and peripheral nerves (**B**) expressing Dicer-2 (Dcr2) alone with robust Sdc expressed in the glial layers (arrows).

(**C,D**) Sdc RNAi-1 expressed with Dicer2 in brain lobe (**A**) and peripheral nerves (**B**) showing loss of Sdc expression in the glial layers (arrows).

(**E**) Cartoon of the *Sdc* gene. Non-coding exons are in gray, coding exons are coloured with extracellular coding exons in dark blue, the conserved Sdc domain in red and intracellular plus transmembrane domain in light blue. The location of the GFP min-exon, the RNAi target sequences and the *Sdc* mutations utilized in this study are indicated.

(**F**) Quantification of Sdc RNAi and Sdc mutant adult viability. Sdc RNAi-1 and RNAi-2 were expressed with Dicer2 (Dcr2) in all glial using repo-Gal4. Knockdown of Sdc lead to a decrease in the number of adults eclosing compared to control (Dcr2). *Sdc* homozygous mutants (*Df(2R)48, ubi-Sara/Sdc*$^{97}$) had reduced viability when compared to heterozygous mutants (*Df(2R)48, ubi-Sara/*+) and controls (*w*$^{1118}$).

### Supplemental Figure 1

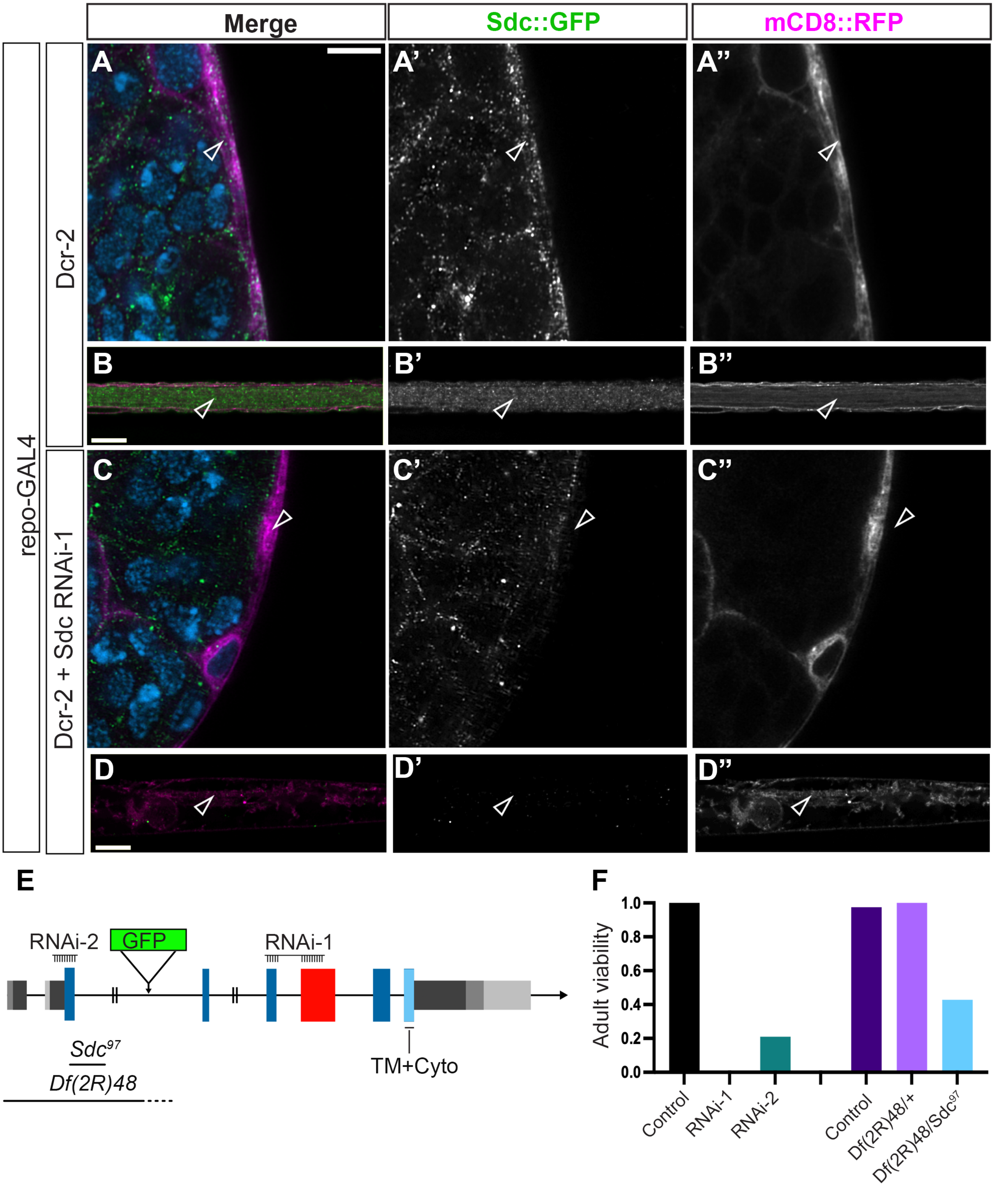
